# Estimating protein isoform abundances with PAQu

**DOI:** 10.64898/2026.04.20.719668

**Authors:** Lorenzo Testa, Lambertus Klei, Alesia Rengle, Anastasia Yocum, David A. Lewis, Bernie Devlin, Kathryn Roeder, Matthew L. MacDonald

## Abstract

A single gene can encode multiple versions of a protein, dubbed isoforms, with varying functionality. Cellular control of isoform abundances is critical for multiple aspects of biology and is only partially regulated by transcript levels. While long-read sequencing facilitates transcript quantification, quantifying the resulting protein isoforms on a large scale is a major challenge, complicating biological interpretation of transcript alterations. Standard “bottom up” mass spectrometry can assess only short portions of isoforms called peptides, and these peptides often map onto more than one isoform. We introduce `PAQu`, a novel Bayesian method that leverages multiomic information from the peptidome and transcriptome to provide accurate estimates of isoform abundance even when peptide mapping is ambiguous. `PAQu` offers several advantages over existing methods in a unified framework. It provides uncertainty quantification, integrates multiomic information for improved accuracy, and provides a rigorous framework for hypothesis testing. Extensive simulations show that `PAQu` consistently outperforms competing methods in detecting differentially expressed protein isoforms and estimating their abundances. We use `PAQu` to investigate differences in isoform abundance levels between people with schizophrenia and control subjects, confirming a long held hypothesis that levels of the C4A isoform of Complement Component 4 are increased in schizophrenia while C4B is not. These results demonstrate that `PAQu` can identify significant variations in isoform abundance levels not previously possible.

## Introduction

Alternative splicing is a crucial post-transcriptional process (*1, 2*) that generates multiple protein isoforms from a single gene, thereby contributing to the complexity of eukaryotic organisms. Dysregulation of this process has been linked to various diseases, including cancer (*3, 4*), neurodegenerative and psychiatric disorders (*5*–*7*), and developmental abnormalities (*8–10*). Despite the significant role of alternative splicing in cellular function (*2*), accurate and comprehensive quantification of protein isoforms remains a major challenge in proteomics (*11, 12*).

In a typical shotgun proteomics experiment, proteins are first digested into peptides and then analyzed by liquid chromatography tandem mass spectrometry (LC-MS/MS). The resulting spectra are used to identify and quantify peptides in digested samples. Although this approach is effective for analyzing individual peptides, it often falls short when the goal is to characterize protein isoforms (*13*). In particular, a significant challenge in protein isoform identification and quantification is the presence of *shared* peptides (*11–13*), which can map onto multiple isoforms of the same protein or even to different proteins due to sequence similarity. Additionally, peptides unique to individual isoforms are typically less abundant compared to those that are shared.

To overcome this challenge, standard practices either collapse proteins with shared peptides into protein groups or ignore shared peptides altogether, which limit isoform characterization (*14–16*). Several other strategies aggregate peptide abundances into protein isoform estimates by taking summary statistics (e.g., average, sum, maximum) without considering whether peptides are shared or isoform-specific. These approaches are reflected in the default settings in the commonly used informatics programs, such as Proteome Discoverer (*17*) and MS Fragger (*18*). As a consequence, standard methods often struggle to distinguish between isoforms with high sequence similarity, leading to underestimation of isoform diversity or potential misinterpretation of biological processes.

To improve isoform quantification, recent approaches have attempted to integrate transcriptomic data with proteomic data, using the relationship between gene expression and protein production. These methods offer a promising direction for tackling the challenge of protein quantification. In particular, several studies such (*19–21*) have introduced techniques that use transcript expression levels as a prior source of information to estimate isoform abundances. However, despite their advances, these methods come with several limitations. For example, Carlyle et al. (*19*) employ an expectation-maximization (EM) algorithm that does not provide any direct measure of uncertainty quantification. Miller et al. (*20*) rely on long-read RNA-seq, an expensive technique to obtain transcript data. Bollon et al. (*21*), exploiting the conjugacy between Dirichlet and Multinomial distributions to accelerate computation, need to discretize the data, resulting in reduced precision. Moreover, none of these methods provides a direct way to evaluate the presence of differentially abundant protein isoforms between different groups of samples.

To overcome these shortcomings, we introduce a new method – which we call `PAQu` – for accurate Protein isoform Abundance Quantification. `PAQu` combines the strengths of transcriptomic and proteomic data, exploiting complementary insights from RNA-Seq and mass spectrometry. It offers robust uncertainty quantification for all estimated parameters, integrates external covariate information, elucidates the relationship between the transcriptome and the proteome, and enables simultaneous differential analysis, providing a comprehensive solution for isoform quantification. Complete `PAQu` code is available on `GitHub` at: https://github.com/testalorenzo/PAQu.

## Results

### Overview of PAQu

`PAQu` is a Bayesian supervised factor analysis method designed for Protein isoform Abundance Quantification. It models protein isoform abundances as latent factors and estimates them by learning two key mappings: one from transcript expression levels to protein isoform abundances, and another from protein isoform to peptide abundances. Simultaneously, `PAQu` assesses the impact of binary conditions on isoform abundances, enabling analysis of differential protein abundance. Our method takes four inputs: a transcript expression matrix from tissue samples from *n* subjects, a peptide abundance matrix from tissue from the same (or possibly different) subjects, a detectability mask matrix indicating whether a peptide *k* can come from a protein isoform *j*, and a binary vector representing a condition for subjects, such as diagnosis (Fig. 1). `PAQu` assumes that transcript expression levels are proportional to the corresponding protein isoform abundances, and that peptide abundances are related to their respective protein isoforms. This naturally leads to a two-layer model. In the first layer, the transcript expression matrix **T** is transformed into the latent protein isoform matrix **I** through a diagonal weight matrix **W**.

**Figure 1:**
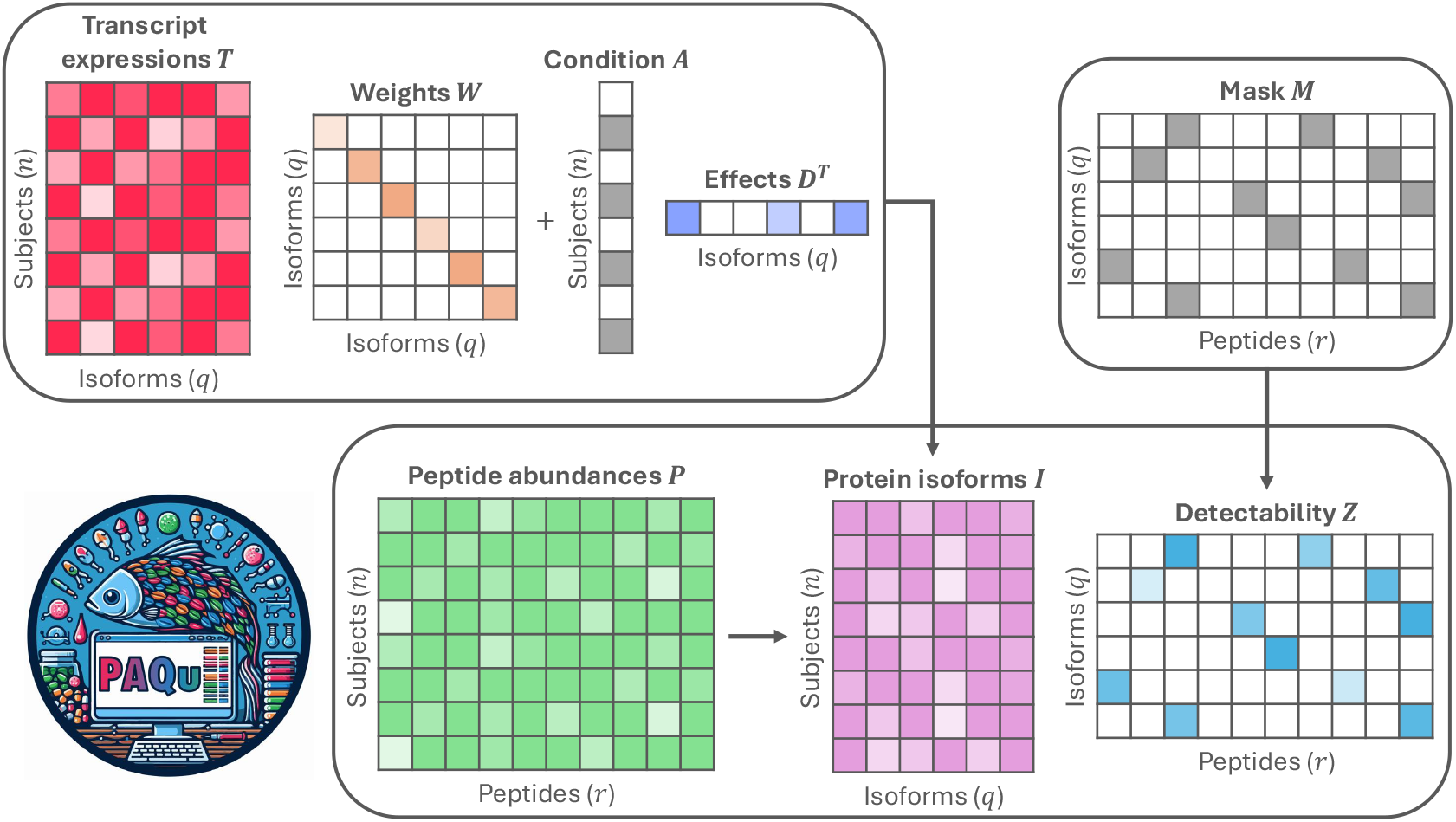
Summary of PAQu model. The model takes as input a transcript expression matrix **T**, a condition binary vector **A**, a detectability mask matrix **M**, and a peptide abundance matrix **P**. The output includes the effects of transcripts on protein isoforms **W**, the difference in protein isoform abundances across conditions **D**, and the estimated associations between isoforms and peptides **Z**.

In the second layer, the peptide abundance matrix **P** is modeled as the product of the protein isoform matrix **I** and the detectability matrix **Z** (whose entries can be zero according to the detectability mask matrix provided as input). Moreover, for differential protein abundance analysis, the first layer also incorporates the relationship between the binary condition vector **A** and the transcript expression matrix **T** through the vector of effects **D**. Conceptually, the first layer performs multivariate linear regression of protein isoforms **I** on transcripts **T** and condition **A**, while the second layer applies factor analysis to **P**. By constraining, in the first layer, the direction of the protein isoform factors within the space spanned by the transcripts, `PAQu` supervises, in the second layer, the estimation of the protein factors. Finally, if available, `PAQu` can also take advantage of the information coming from external covariates, easily inserting them into the previous model (see Methods).

Although this two-layer problem is not identifiable in the traditional frequentist sense, imposing prior distributions on the unknowns allows estimation of the quantities of interest. Specifically, for the `PAQu` model, we use Gaussian priors for the protein isoform abundances **I**, the weights in **W**, the effects in **D**, and truncated Gaussian priors for the detectability factors in **Z**. For situations with a high signal-to-noise ratio, users can opt for a sparser “spike-and-slab” prior (*22*), which offers more conservative inference by selectively shrinking smaller effects.

We implement a Gibbs sampling algorithm to generate posterior samples of the model parameters (*23*). The significance of any effect can be measured by the local false sign rate (LFSR) metric (*24*), a summary of the posterior distribution akin to local false discovery rate (see Methods). Similarly, for parameters with sparse priors, posterior inclusion probability (PIP) can be easily computed.

`PAQu` generates several outputs: estimated associations between transcripts and protein isoforms, inferred associations between protein isoforms and peptides, estimated abundances of protein isoforms, and a list of differentially abundant protein isoforms, which can be interpreted, for instance, through gene ontology (GO) enrichment analysis of genes associated with each isoform.

All of these estimated quantities are accompanied by a measure of their uncertainty.

### Simulation results provide strong motivation for PAQu

We evaluate the performance of `PAQu` in two main simulation settings. In the first setting, which we refer to as “easy”, we generate data without shared peptides. Specifically, we sample one transcript (*q* = 1) from a Normal distribution, map it into a protein isoform with random conversion weight, add an effect due to a binary condition, and then split it into two peptides (*r* = 2) using randomly assigned detectability scores (see Methods). This setting serves as a reference point to compare `PAQu` with conventional methods for protein quantification. We run each simulation experiment with three different sample sizes (*n* = 100, 200, 500), three effect sizes for the binary condition (**D** _*j*_ = 0.33, 0.66, 1), and with 25 different seeds. In the second, more challenging scenario – referred to as the “difficult” setting – we generate data that include shared peptides. Here, we sample five transcripts (*q* = 5) from a multivariate Normal distribution, map them to their corresponding protein isoforms with random conversion weights, and introduce a binary condition effect to some isoforms. These isoforms are split into 10 peptides (*r* = 10) using random detectability scores, with 30% of the detectability matrix entries being nonzero. Each simulation is run across three sample sizes (*n* = 100, 200, 500), three effect sizes (**D** _*j*_ = 0.33, 0.66, 1), three numbers of differentially abundant isoforms (|**D**^*act*^ | = 1, 2, 3) and 25 different random seeds.

We exploit simulated data to assess the performance of various prior distributions in differential isoform abundance tests. In our simulations, the effects inferred by using the “spike-and-slab” prior are overly conservative, particularly when the signal or the sample size is small. In contrast, the Normal prior exhibits much less bias (*SI Appendix*, Fig. 6 and 7), which supports our decision to adopt the Normal prior as default. We then evaluate the ability of `PAQu` to recover and quantify differentially abundant protein isoforms. `PAQu` accurately estimates these effects, showing a slightly conservative behavior in the simulation setting “hard” in terms of recovery of differentially abundant isoforms at LFSR ≤0.05 when the sample size and the signal are small (*SI Appendix*, Fig. 8 and 9). This leads to a modest increase in false negatives, as expected due to the conservative prior used and the extreme weakness of the signal.

To assess the performance of `PAQu` in inferring the latent protein isoform factors, we measure the absolute correlation between the inferred and true isoform abundances. In all scenarios, the estimated protein factors show a strong correlation with the true factors (Fig. 2). As expected, the quality of the estimates increases with the sample size; at the same time, estimates tend to slightly worsen with increasing number of parameters to estimate.

**Figure 2:**
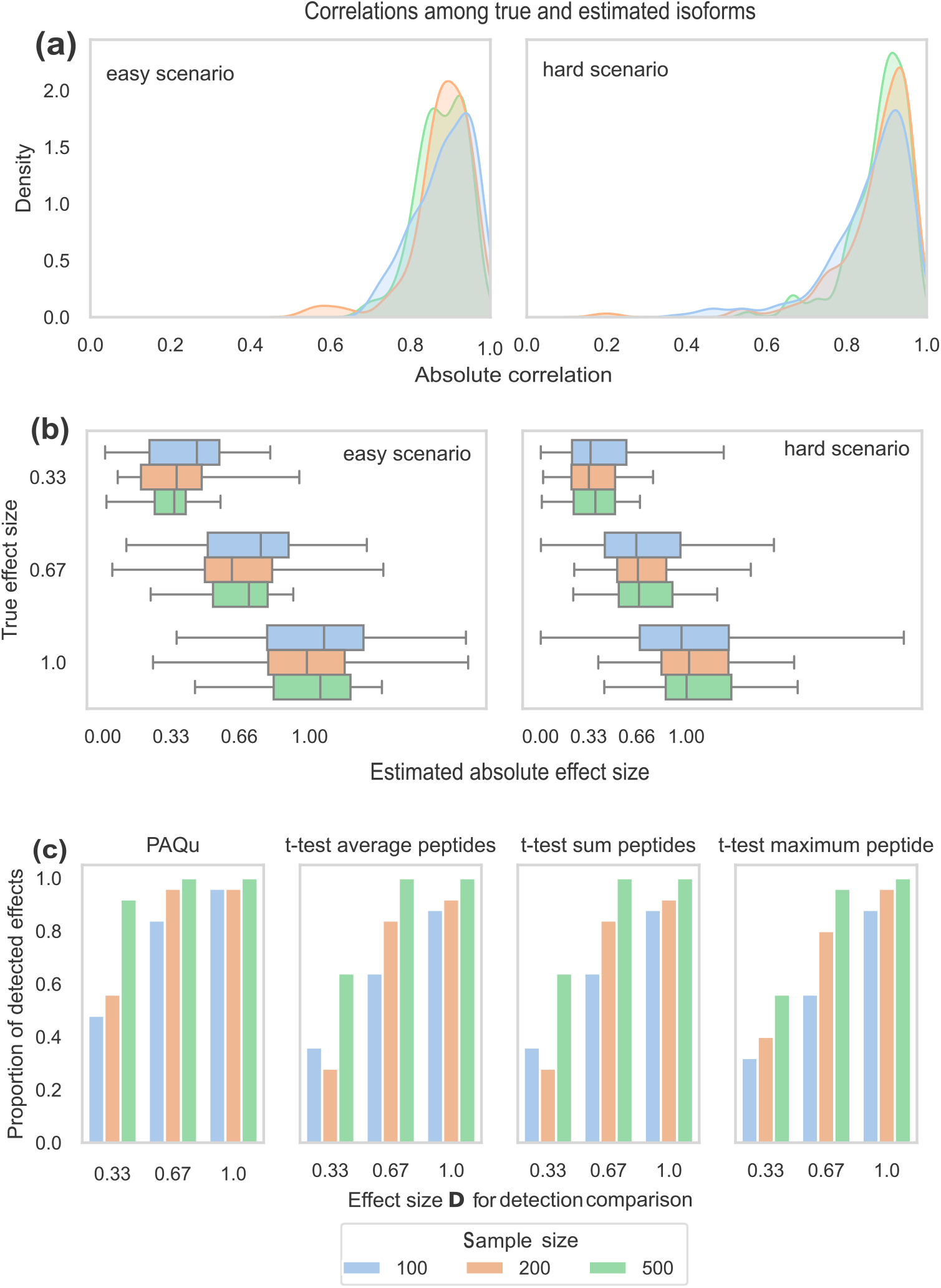
Simulation results. **(a)** Distributions of the absolute correlation values between true protein isoforms and the estimates inferred by `PAQu` under the “easy” (left) and “difficult” (right) scenarios. Different colors represent different sample sizes, ranging from 100 to 500. **(b)** Box plots of absolute effect sizes estimated by `PAQu` under the “easy” (left) and “difficult” (right) scenario. Different colors represent different sample sizes. For each box, the center line represents the median; the lower and upper hinges correspond to the first and third quartiles; the upper and lower whiskers span 2 times the interquartile range. **(c)** Comparison between the proportion of detected effects **D** _*j*_ under the “easy” scenario between `PAQu` (LFSR ≤0.05), t-test on the average of peptide abundances, t-test on the sum of peptide abundances, t-test on the maximum peptide abundance (p-value ≤ 0.05). All simulations are repeated 25 times.

We then compare the performance of `PAQu` in detecting differentially abundant isoforms with other commonly used tools: t-test on the average abundances of peptides, t-test on the sum of peptides, and t-test on the maximum peptide abundance. `PAQu` outperforms all these methods in all simulation settings (Fig. 2, *SI Appendix*, 8 and 9). These results demonstrate not only the performance gains `PAQu` can provide, but also the value of integrating data from both the transcriptome and the peptidome to achieve more precise isoform-level estimates.

Moreover, to further challenge `PAQu` and measure its performance under misspecification of the model, we evaluate simulations in which we assume that there is no link between transcript expression levels and protein isoform abundances (i.e. **W** = 0). Remarkably, even when transcripts carry no information about protein isoforms, `PAQu` can still provide meaningful inference, significantly outperforming competitors (*SI Appendix*, Fig. 10, 11, and 12). To evaluate the contribution of information from transcript expressions, we performed simulations in which we compared the estimates of `PAQu` without transcripts, only average values between samples and full transcript expression levels. As the amount of transcript expression increases, so does the quality of isoform abundance estimates, measured by the absolute correlation between true and estimated abundances (*SI Appendix*, Fig. 13). Finally, we assess computational feasibility. Although *T* and *P* can be large, because *Z* is block diagonal, computations via the Gibbs sampler can proceed in parallel by block, making them efficient (*SI Appendix*, Fig. 14).

### PAQu reveals the effects of schizophrenia risk genes

We applied `PAQu` to the most deeply characterized proteomic dataset of human brain tissue for a psychiatric disorder. Quantitative proteomics using TMT labeling and deep offline fractionation was applied to postmortem dorsal anterior cingulate cortex gray matter from 56 individuals diagnosed with schizophrenia and 56 matched controls with no history of psychiatric or neurological conditions. While “protein” expression alterations have previously been observed in schizophrenia, here we report differentially abundant protein isoforms. Furthermore, we have complementary data on transcripts, based on RNA-seq measurements (*25*), in adjacent tissue sections from a subset of the same subjects. After collecting and preprocessing data (see Methods), we have protein and transcript data from *n* = 103 subjects, 53 of whom are affected by schizophrenia, and consisting of *q* = 9169 isoforms and *r* = 79, 040 peptides. Groups of isoforms that shared exactly the same measured peptides, and therefore were not uniquely estimable, were rolled up into isoform groups, which resulted in 1440 isoform groups representing 3338 isoforms and 5831 single peptides. The isoform groups can be divided into 1396 and 44 groups mapping to a single or multiple genes, respectively. `PAQu` calculations were performed using the combined 7271 single and grouped isoforms.

We ran `PAQu` with 10 different MCMC chains, each consisting of 3000 iterations (the first 2000 are burn-in), both to measure `PAQu` consistency and to gauge the stability of our findings. For differential case-control abundance, the *SI Appendix*, Fig. 15 shows the detection frequency results across chains at LFSR ≤0.05. Out of 7271 isoforms, `PAQu` never selects 6211 isoforms (85.4%) as differentially abundant, and always detects 541 isoforms (7.4%). The remaining 519 (7.1%) isoforms are selected at least once, but only 179 of them are detected more than 5 times. The high degree of consistency among runs further strengthens confidence in the interpretation of the findings obtained by using `PAQu`.

We group isoforms on the basis of the number of times they are selected across MCMC chains. Then, for each level of minimum agreement factor *γ* among `PAQu` runs, we map protein isoforms back to their associated genes, and then perform Gene Ontology (GO) enrichment analysis (Fig. 3). Notably, for more lenient thresholds for significance, synaptic and neuronal biology emerge from the gene set, whereas themes involving energetics (mitochondrial) and protein synthesis (ribosomal) are always over-represented.

**Figure 3:**
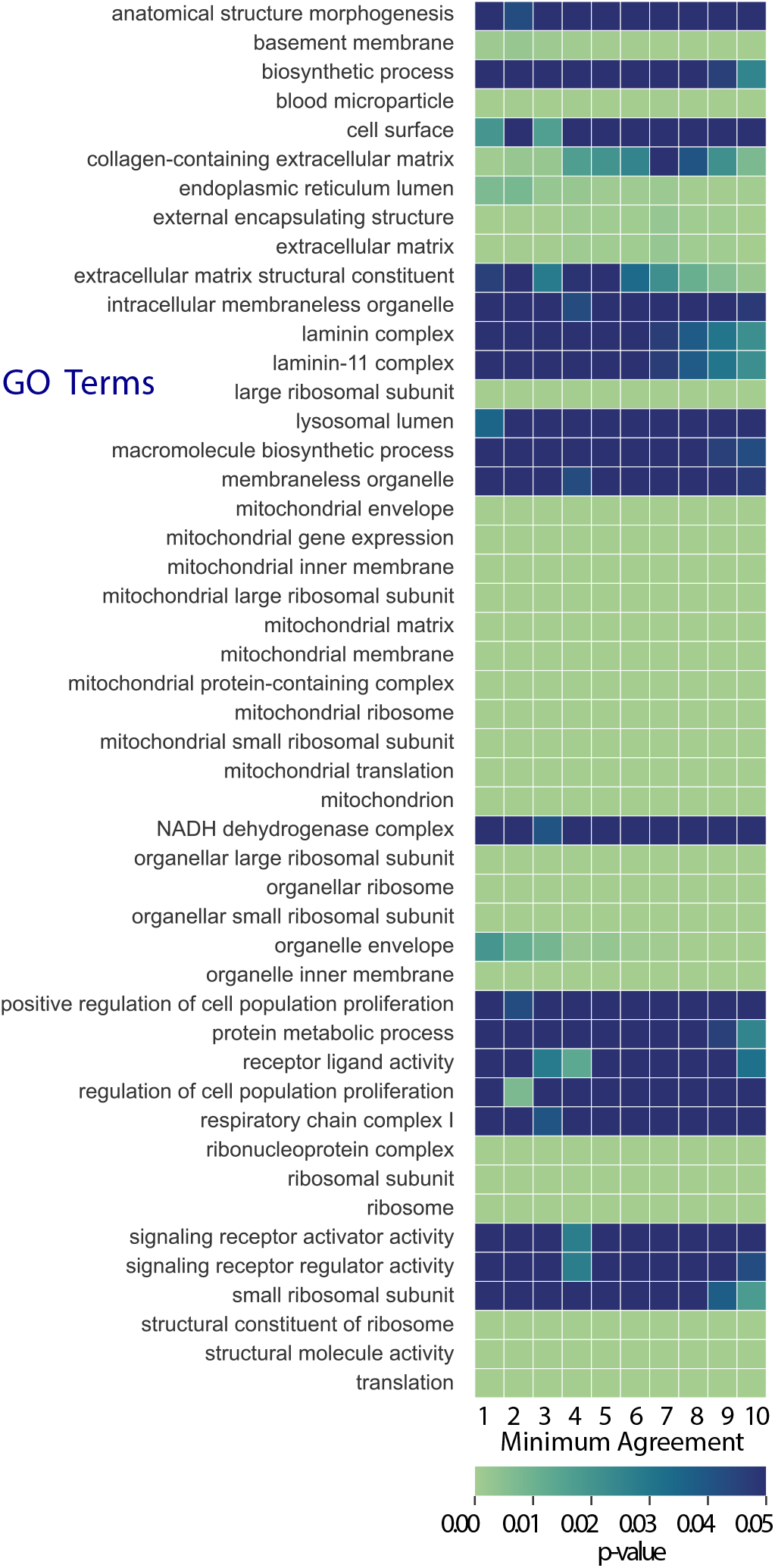
GO enrichment analysis. Isoforms are characterized by the number of times they are selected across `PAQu` MCMC chains. Then, for each level of minimum agreement factor *γ* among `PAQu` runs, we map protein isoforms meeting this criterion back to their associated genes and then perform Gene Ontology (GO) enrichment analysis.

### PAQu sheds light on the relationship between transcriptome, proteome, and peptidome

`PAQu` also provides estimates of the links between transcripts and protein isoforms. Using the same data as described above, we analyze the posterior distributions of the conversion coefficients **W** (Fig. 1). Interestingly, the distribution of the coefficients related to transcripts and protein isoforms, which are detected as significant in their respective domains, is shifted rightward (Fig. 4A,B). This suggests that in the presence of biological variation in both the transcriptome and the proteome, statistical relationships arise naturally. We decided to investigate this further because the relationship between gene expression measured in tissue and the amount of corresponding protein found in the same tissue has been an intriguing and important scientific puzzle (*26, 27*). In some experimental settings, gene expression and protein abundance are highly correlated (*28–30*); in survey studies, the correlation is often low, including most studies evaluating brain tissue (*31, 32*). While our analyses bring the scale down to RNA transcripts and their protein isoforms, the parallels to previous work are obvious.

**Figure 4:**
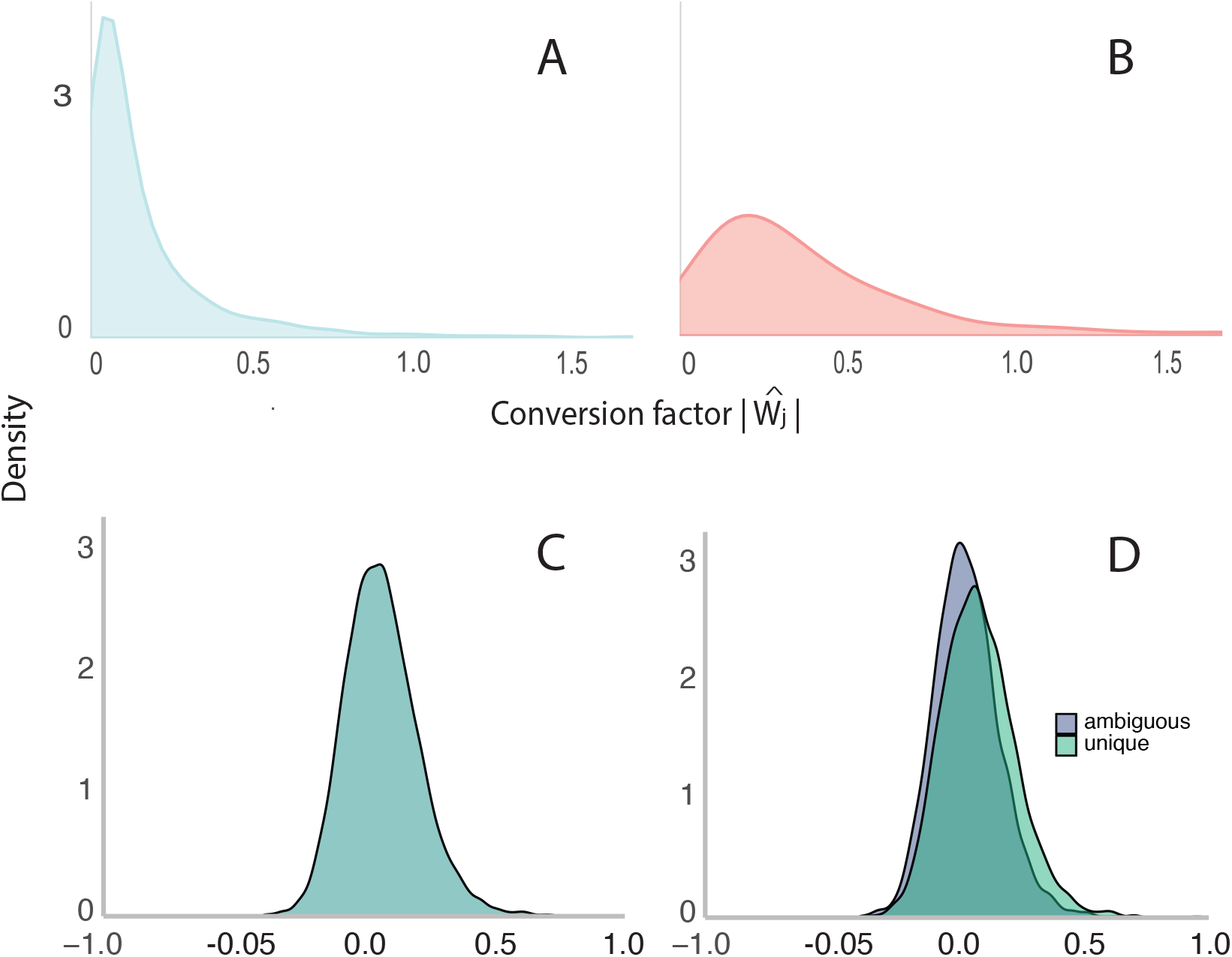
Relationship of transcripts and protein isoform abundance within individuals. (A-B) Transcripts and protein isoforms that show biological variation (*γ* > 5): (A) for all, (B) detected by t-test on transcripts and PAQu. (C) Correlation between all transcripts and their isoform abundances; and (D) Correlation between transcript and isoform abundance split by whether there is a unique or ambiguous mapping of peptides onto transcripts/isoforms.

Of the 7271 isoforms, 7107 show limited variability between MCMC chains (Methods). For each of these reliably estimated isoforms, we computed the correlation of isoform and transcript abundances over individuals. We obtain a roughly symmetric distribution centered at 0.067 but showing right skew (Fig. 4C). If the true transcript/isoform correlation were zero, then the observed distribution would be normally distributed about zero and only 0.2% would exceed 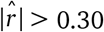, about 7 observations exceeding each bound. Instead, we found 417 transcripts/isoforms with correlations > 0.3, 5.9% of the pairs. We also observed 19 transcripts/isoforms with unusually low correlations, below − 0.3 (0.3%). Note that isoforms can be split into two distinct categories: isoforms with a unique mapping of peptides onto isoform (unique) and those for which the measured abundance of a peptide could trace to two or more isoforms (ambiguous).

For the 4056 unique transcripts /isoforms (Fig. 4D), their distribution is centered at 0.087, with 316 (7.8%) in the positive tail and only 6 (0.1%) strongly negative, as previously defined. For the 3051 in the ambiguous set (Fig. 4D), the distribution is centered at 0.041 and it has 101 (3.3%) in the positive tail and 13 (0.4%) in the negative tail. Thus, when there is greater uncertainty in how peptide abundances inform isoform abundances, `PAQu` is somewhat less accurate, leading to fewer strongly positive correlations.

Next, we focus on extreme positive correlations. As we develop in *SI Appendix*, Table 1, these correlations must arise from substantial biological variation among samples for both transcripts and isoforms. For example, if cases and controls differed in their means by one half of a standard deviation for both transcript and complementary isoform – an unusually large difference – the expected correlation induced would only be *r* ≈ 0.053. To determine what variation predicts the observed correlation for each (isoform, transcript) pair, we build a model using the following five predictors: mean case-control difference for isoform/transcript abundances; coefficient for the effect of age on isoform/transcript abundances; and impact of genetic variation on the gene. For this last predictor, we use results from GTEx (*33*) analysis of gene expression. Specifically, we use the variation of gene expression explained by the eGENE genetic variant (Methods). Using these predictors, we fit the model for two classes of outcomes: the 412 (transcript,isoform) pairs with strong positive correlation values, *r* > 0.3 (5 others did not map uniquely to a single gene); and 412 randomly-chosen (transcript,isoform) pairs, conditional on their correlation being in the range of |*r* | < 0.05. Using all 824 outcomes, the model is significant, with *R*^2^ = 0.061 (*SI Appendix*, Table 1). The impact of age on transcript abundance, diagnosis-based mean isoform differences, and eGENE variation explain a significant portion of the variation. If the low correlation set is analyzed, no predictor explains a significant amount of the variation; when the strong positive correlation set is analyzed, age and diagnosis for both transcripts and isoforms are important predictors, whereas genetic variation is not. The model accounts for a slightly larger portion of the variability of these higher correlation outcomes (*R*^2^ = 0.092); *SI Appendix*, Table 1).

**Table 1:** Results from linear model analysis on three groups: both high and null correlations, only null correlations (|*r*| < 0.05), and high correlations (*r* > 0.3).

| Predictors<sup>1</sup> | Estimate | Std. Error | t value | P-value |
| --- | --- | --- | --- | --- |
| intercept | 0.184 | 0.008 | 23.574 | 0.000 |
| diagnosis, isoform | 0.225 | 0.080 | 2.808 | 0.005 |
| age, isoform | -0.135 | 1.549 | -0.087 | 0.931 |
| diagnosis, transcript | 0.035 | 0.049 | 0.716 | 0.474 |
| age, transcript | 4.831 | 1.325 | 3.646 | 0.000 |
| genetic variation | 0.370 | 0.126 | 2.938 | 0.003 |

**Null correlation group**
| Predictors<sup>1</sup> | Estimate | Std. Error | t value | P-value |
| --- | --- | --- | --- | --- |
| intercept | 0.003 | 0.002 | 2.066 | 0.039 |
| diagnosis, isoform | 0.030 | 0.021 | 1.411 | 0.159 |
| age, isoform | -0.450 | 0.418 | -1.077 | 0.282 |
| diagnosis, transcript | -0.001 | 0.011 | -0.101 | 0.920 |
| age,transcript | -0.437 | 0.277 | -1.576 | 0.116 |
| genetic variation | -0.012 | 0.034 | -0.362 | 0.717 |

**High correlation group<sup>2</sup>**
| Predictors<sup>1</sup> | Estimate | Std. Error | t value | P-value |
| --- | --- | --- | --- | --- |
| intercept | 0.388 | 0.005 | 76.082 | 0.000 |
| diagnosis, isoform | 0.246 | 0.045 | 5.457 | 0.000 |
| age, isoform | -2.936 | 0.940 | -3.125 | 0.002 |
| diagnosis, transcript | -0.058 | 0.030 | -1.917 | 0.056 |
| age, transcript | 2.220 | 0.898 | 2.471 | 0.014 |
| genetic variation | 0.103 | 0.066 | 1.559 | 0.120 |

<sup>1</sup>All predictors are estimated effects from analysis of the specific data type.

Earlier research (*31*) found that gene expression within certain gene sets shows higher correlation to resulting protein abundance. To determine if our results show a similar pattern, we next performed a gene ontology overrepresentation analysis (*34–36*), with the universe set to the genes observed in our data. We queried cellular components (CC) and biological processes (BP). For the genes mapping onto the high correlation set, 40 gene sets show significant enrichment (FDR < 0.05) (*SI Appendix*, Table 2). Comparing our enriched gene sets with the thirty significantly enriched gene sets from (*31*), two gene sets overlapped, namely cell surface and extracellular matrix. Given the large set of possible terms for CC and BP, two overlapping gene sets is larger than that expected by chance, and it is perhaps more remarkable because (*31*) took a very different analytical approach to assess a somewhat different data type.

**Table 2:** GO Cellular Component and Biological Process Analysis.

| GO cellular component | Genes in set | Observed genes | Expected | Fold Enrichment | FDR | GO Type |
| --- | --- | --- | --- | --- | --- | --- |
| actin-based cell projection (GO:0098858) | 97 | 19 | 6.95 | 2.74 | 0.00371 | CC |
| adherens junction (GO:0005912) | 112 | 18 | 8.02 | 2.24 | 0.0463 | CC |
| apical part of cell (GO:0045177) | 177 | 25 | 12.67 | 1.97 | 0.0398 | CC |
| astrocyte projection (GO:0097449) | 16 | 6 | 1.15 | 5.24 | 0.0301 | CC |
| axon (GO:0030424) | 447 | 52 | 32.01 | 1.62 | 0.0268 | CC |
| blood microparticle (GO:0072562) | 35 | 13 | 2.51 | 5.19 | 0.0000997 | CC |
| brush border (GO:0005903) | 44 | 12 | 3.15 | 3.81 | 0.00355 | CC |
| cell junction (GO:0030054) | 1467 | 146 | 105.04 | 1.39 | 0.000596 | CC |
| cell periphery (GO:0071944) | 1969 | 195 | 140.98 | 1.38 | 0.00000713 | CC |
| cell surface (GO:0009986) | 265 | 40 | 18.97 | 2.11 | 0.000619 | CC |
| cluster of actin-based cell projections (GO:0098862) | 61 | 17 | 4.37 | 3.89 | 0.000161 | CC |
| external encapsulating structure (GO:0030312) | 115 | 27 | 8.23 | 3.28 | 0.00000849 | CC |
| extracellular matrix (GO:0031012) | 115 | 27 | 8.23 | 3.28 | 0.00000679 | CC |
| extracellular region (GO:0005576) | 1436 | 141 | 102.82 | 1.37 | 0.00127 | CC |
| extracellular space (GO:0005615) | 1272 | 128 | 91.08 | 1.41 | 0.00132 | CC |
| intermediate filament (GO:0005882) | 29 | 8 | 2.08 | 3.85 | 0.0354 | CC |
| intermediate filament cytoskeleton (GO:0045111) | 34 | 9 | 2.43 | 3.70 | 0.0272 | CC |
| neuron projection (GO:0043005) | 785 | 81 | 56.21 | 1.44 | 0.0272 | CC |
| plasma membrane (GO:0005886) | 1842 | 172 | 131.89 | 1.30 | 0.00185 | CC |
| presynapse (GO:0098793) | 474 | 53 | 33.94 | 1.56 | 0.0357 | CC |
| synapse (GO:0045202) | 1164 | 110 | 83.34 | 1.32 | 0.0459 | CC |
| synaptobrevin 2-SNAP-25-syntaxin-1a complex (GO:0070044) | 7 | 4 | 0.5 | 7.98 | 0.0361 | CC |
| anterograde trans-synaptic signaling (GO:0098916) | 226 | 34 | 16.18 | 2.1 | 0.0141 | BP |
| carbon dioxide transport (GO:0015670) | 4 | 4 | 0.29 | 13.97 | 0.0137 | BP |
| cell adhesion (GO:0007155) | 362 | 51 | 25.92 | 1.97 | 0.00523 | BP |
| cell-cell signaling (GO:0007267) | 343 | 48 | 24.56 | 1.95 | 0.00389 | BP |
| chemical synaptic transmission (GO:0007268) | 226 | 34 | 16.18 | 2.1 | 0.0147 | BP |
| kidney epithelium development (GO:0072073) | 34 | 10 | 2.43 | 4.11 | 0.0313 | BP |
| modulation of chemical synaptic transmission (GO:0050804) | 327 | 47 | 23.41 | 2.01 | 0.00389 | BP |
| multicellular organismal process (GO:0032501) | 2042 | 185 | 146.21 | 1.27 | 0.0205 | BP |
| nervous system process (GO:0050877) | 364 | 50 | 26.06 | 1.92 | 0.00396 | BP |
| one-carbon compound transport (GO:0019755) | 5 | 4 | 0.36 | 11.17 | 0.0412 | BP |
| regulated exocytosis (GO:0045055) | 82 | 18 | 5.87 | 3.07 | 0.00848 | BP |
| regulation of ERK1 and ERK2 cascade (GO:0070372) | 94 | 20 | 6.73 | 2.97 | 0.006 | BP |
| regulation of synaptic transmission glutamatergic (GO:0051966) | 41 | 11 | 2.94 | 3.75 | 0.0342 | BP |
| regulation of trans-synaptic signaling (GO:0099177) | 328 | 47 | 23.49 | 2 | 0.00364 | BP |
| synaptic signaling (GO:0099536) | 267 | 39 | 19.12 | 2.04 | 0.00827 | BP |
| synaptic vesicle docking (GO:0016081) | 13 | 6 | 0.93 | 6.45 | 0.0459 | BP |
| system process (GO:0003008) | 642 | 75 | 45.97 | 1.63 | 0.00692 | BP |
| trans-synaptic signaling (GO:0099537) | 245 | 37 | 17.54 | 2.11 | 0.00694 | BP |

We noted a complementary feature of the genes encoding transcripts/isoforms in the high correlation set, namely they were highly enriched for terms related to neuronal and synaptic biology. We used ReviGO (*37*) to summarize these terms into coherent themes (Fig. 5). This result is consistent with prior studies that show tight scaling of transcription and translation based on synaptic activity (*38, 39*), although this tight coupling potentially diminishes with increasing age (*40, 41*).

**Figure 5:**
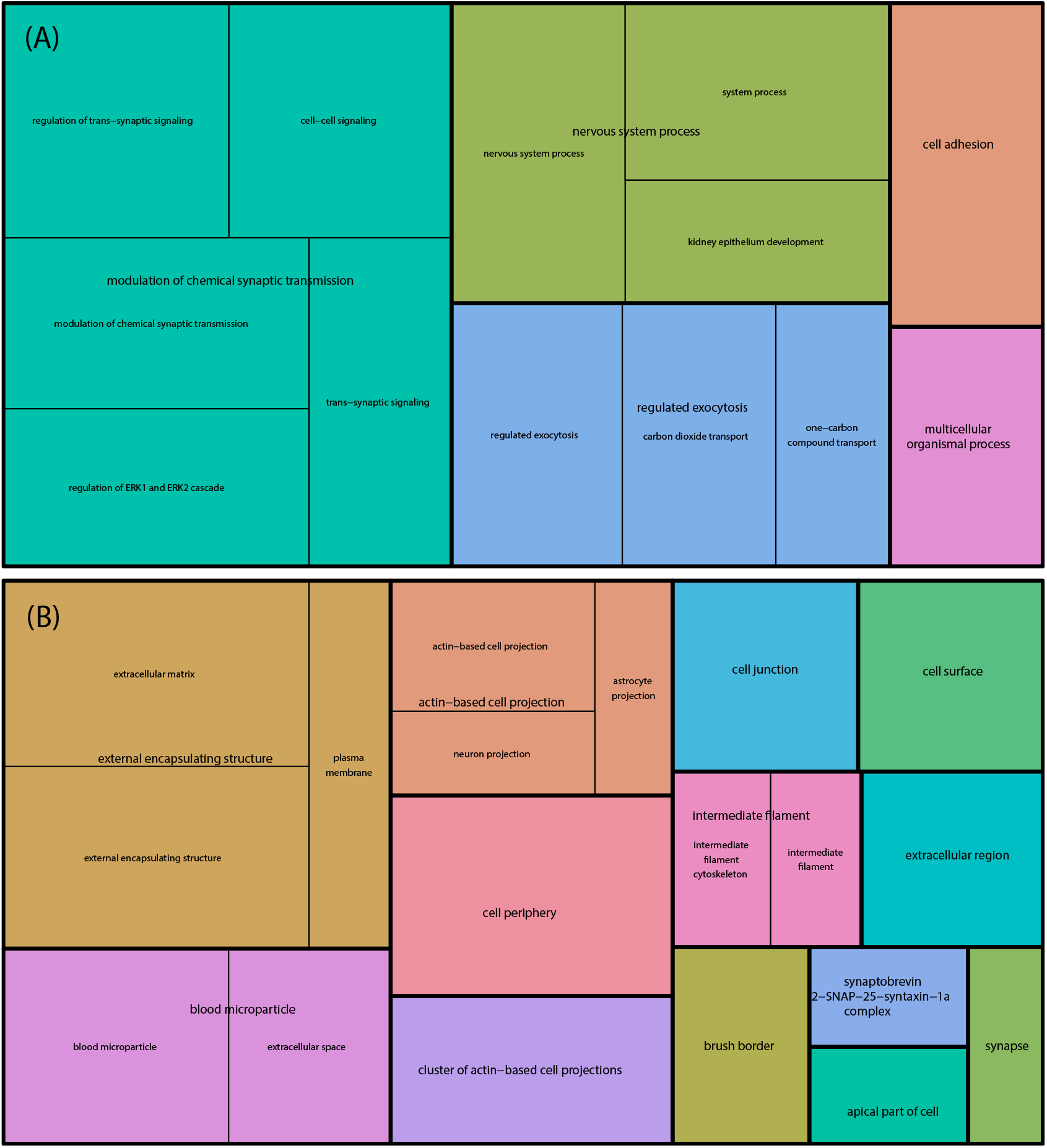
Revigo tree plot summary of enriched Gene Ontology terms for transcripts and isoforms showing large correlaton of abundances. (A) Term enrichment for Biological Processes; (B) Term enrichment for Cellular Component.

`PAQu` also returns estimates of the detectability matrix **Z**, which projects peptide abundances onto isoform abundances. Peptides vary in their detectability by the mass spectrometer and spectral matching algorithms for a host of reasons, such as their amino acid composition and how those amino acids are chemically altered in vivo, i.e., post-translational modifications (PTMs). For the latter, while the peptide itself has the same amino acid sequence, PTMs change its mass/charge signature and thus the detectability as the same peptide (*42*). In turn, PTMs can induce variability in measured abundances of peptides from the same isoform and their relationship to the isoform.

## Discussion

Advanced transcriptomics approaches facilitate quantification of transcript alterations in disorders like schizophrenia that could contribute to disease pathology. However, confirming these observations at the isoform level has proven challenging due to bottle necks in proteomic data acquisition and analysis. `PAQu` stands as an effective tool for addressing several of these challenges: it provides uncertainty quantification for all estimated parameters, integrates external covariate information, elucidates the relationship between the transcriptome and the proteome, highlights the potential presence of post-translational modifications, and allows users to perform differential abundance analysis.

Our analyses of transcriptome and proteome from postmortem dorsal anterior cingulate cortex (dACC) gray matter reveal close integration of certain neuronal and synaptic transcripts and isoforms (Fig. 4, Fig. 5). Meaningful biological variation, including changes in abundance with aging, based on case-status, and genetic variation determine some of this integration (Table 1). For instance, among the 412 highly correlated transcripts and isoforms (*r* > 0.3), 47 are classified under the term regulation of trans-synaptic signaling (GO:0099177), and these show the greatest enrichment for Gene Ontology’s Biological Processes.

Our results on case-control differences for transcripts and isoforms confirm a critical observation made in schizophrenia, first published by Sekar et al. (*43*) 10 years ago. Elevated levels of the Complement Component 4 transcript C4A, but not C4B, are a genetically based risk factor for schizophrenia, directly correlated to structurally diverse risk alleles. This increase in C4A is widely believed to contribute to impaired cognitive and sensory processing in schizophrenia via its promotion of dendritic spine pruning (*44*). As these isoforms are almost identical, antibodies cannot distinguish between the two, and thus the findings could not be confirmed at the protein level. Our data captures 38 peptides of C4, 36 mapping to both isoforms and 2 mapping exclusively to C4B. Our results show, for the first time, that both C4A transcript and protein isoform levels are significantly increased in schizophrenia. By contrast, and as predicted by transcript levels, the abundance of C4B did not change with case status. We find that several other transcript alterations with relevance for glial and synaptic processes, implicated together in schizophrenia, are realized at the isoform level. For example, CD44 isoform P16070 is also significantly elevated in cases, which is consistent with recent results showing upregulation of reactive astrocyte marker CD44 in the presence of elevated levels of C4A (*45*). However, not all synaptic transcript changes result in concurrent isoform alterations, as is the case in CASKIN2, a key molecule for excitatory synaptic function (*46*). Levels of the CASKIN2 transcript ENST00000321617 are decreased whereas its isoform, Q8WXE0, is significantly elevated, possibly due to a compensatory mechanism to maintain synaptic transmission. Collectively, our findings in schizophrenia demonstrate the power of `PAQu` to provide confirmation that transcript level alterations manifest at the protein level, providing a direct link from genetic risk to protein function while also showing that not all such changes are translated.

`PAQu` has limitations; for instance, it can only work in scenarios when *q* ≤*r*. We envision several possible extensions for our method. First, in situations where the number of samples and the number of isoforms with shared peptides are large, the computational load could be significantly reduced by replacing sampling with a variational inference approach. Second, we would like to investigate the effect of post-translational modifications in greater detail, perhaps by extending `PAQu` to incorporate information regarding them. Third, we would like to extend `PAQu` as to integrate data from additional domains other than the transcriptome and peptidome.

In conclusion, in this paper we propose `PAQu`, a powerful new framework for the analysis of mass spectrometry data that takes advantage of additional information about the transcriptome to quantify the abundance of protein isoforms. By implementing a version of Bayesian supervised factor analysis, we believe that `PAQu` has the potential to help researchers make full use of mass spectrometry data.

## Materials and Methods

### Notation and PAQu model

We define the *n × q* matrix of transcript expression data as **T**, with **T**_*i j*_ representing its (*i, j*)-th element. Similarly, the *n × r* matrix of peptide abundance data is denoted by **P**, where **P**_*ik*_ is the (*i, k*)-th element. The vector **A** is an *n ×* 1 binary indicator describing the condition of sample *i*, with **A**_*i*_ ∈ {0, 1}. The indices *i* = 1, …, *n, j* = 1, …, *q*, and *k* = 1, …, *r* refer to the samples, isoforms, and peptides, respectively. Our objective is to infer the parameters that link these quantities, as described by the following model:

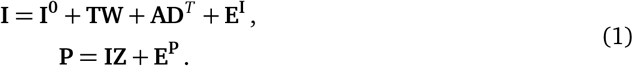

Here, **I** is an *n × q* matrix where **I**_*i j*_ represents the unobserved abundance of isoform *j* in sample *i*, and **I**^**0**^ is an *n × q* matrix of intercepts, with 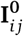 indicating the baseline abundance of isoform *j* across samples (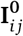 is constant for every *i*). The diagonal matrix **W** has elements **W** _*j j*_, representing the conversion weight between transcript *j* and its corresponding protein isoform. The vector **D** is *q ×* 1, where each **D** _*j*_ describes the effect of the binary condition **A** on the abundance of protein isoform *j*. The matrix **Z** is a *q × r* matrix of detectability coefficients, with **Z**_*jk*_ = 0 if peptide *k* is not associated with isoform *j*, and otherwise unknown. The terms **E**^**I**^ and **E**^**P**^ are random heteroskedastic Gaussian noise terms affecting isoform and peptide abundances, respectively. Note that knowledge of **T** and **A** is not strictly necessary: `PAQu` can operate as long as **P** and a detectability mask are provided. We include here these additional components in the model for the sake of completeness. In *SI Appendix*, Section A, we provide details and in Section B, we describe how to use `PAQu` in cases where information about **T** and **A** is limited or absent, or in the presence of external covariates.

### Estimation and inference with PAQu

#### Prior distributions

The parameters in the model described in Eq. 1 are not identifiable. Thus, standard optimization procedures can lead only to locally optimal solutions. A viable alternative is provided by the Bayesian framework, which requires specification of prior distributions on parameters. We therefore assume that any conversion weight between a transcript *j* and its protein isoform follows a Gaussian distribution:

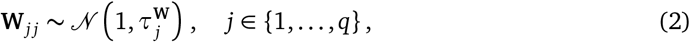

where 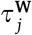 denotes the prior variance. The prior mean is set to 1, which follows the theoretical assumption of perfect proportionality between transcript expression levels and corresponding protein isoform abundances.

We impose a similar Gaussian prior on condition effects **D**:

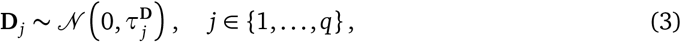

where 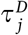 denotes the prior variance, and the prior mean is set to 0 to reflect conservative prior knowledge about potential differential isoform abundances.

The prior distribution for detectability scores depends on the compatibility between protein isoforms and peptides. We write *k* ≺ *j* if a peptide *k* is compatible with an isoform *j*. Therefore, we can write the prior distribution as a mixture between a truncated Gaussian and a Dirac mass at 0:

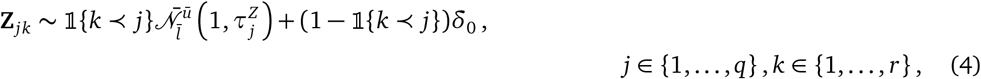

where 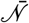 denotes a Truncated Normal distribution with prior mean 1, prior variance 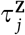, and lower and upper truncation parameters 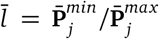 and 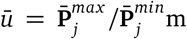 with 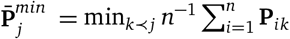 and 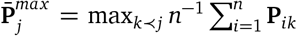. We employ this truncated prior specification for **Z** to guarantee that estimates in **I** are consistent to observed values at the peptide level.

We also specify a Gaussian prior for the intercept **I**^**0**^:

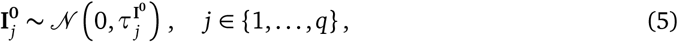

where 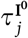 denotes the prior variance. Notice that in an experiment where transcript expression levels have not been measured, but the scientist has access to average expression levels from another study, these may be used as prior means for the intercept (see *SI Appendix*, Section B for details).

In alternative to the Gaussian prior, `PAQu` offers the possibility of implementing a “spike-and-slab” prior. For example, the “spike-and-slab” prior for **D** reads as:

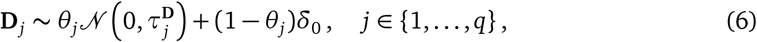

where *θ*_*j*_ is the prior probability that a given isoform *j* is differentially abundant across conditions. As supported by our simulation study, we prefer to use Gaussian priors as they lead to more sensible results.

The last fundamental prior distribution, which links the first and second layers (Fig. 1), is the one for **I**. Again, we assume a Gaussian prior, this time informing it with knowledge about the other covariates:

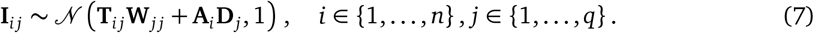

Intuitively, the prior distribution in the previous Equation guarantees that the protein isoforms approximately lie in the space spanned by transcripts. This allows `PAQu` to supervise the factor analysis of peptides in the second layer in a smooth and unified manner. The prior distributions of the other parameters in the model are specified in *SI Appendix*, Section A.

#### Posterior sampling

We approximate the posterior distribution of the model parameters using Gibbs sampling, which is a natural choice since we can derive closed-form expressions for the *full conditionals*. Intuitively, given the values of (**W, D**), **I**, or **Z**, the remaining unknown parameters can be easily determined by linear regression. This approach simplifies the complexity of estimating a multivariate posterior distribution by breaking it down into several manageable univariate problems. The analytical posterior distributions of all parameters in the model – using both Gaussian and “spike-and-slab” priors – are specified in the *SI Appendix*, Section A.

Our choice of default parameters for the Gibbs sampling procedure includes 3000 MCMC iterations and a burn-in period of 2000 iterations. As a result, each MCMC chain approximates its corresponding conditional posterior distribution using 1000 samples.

#### Measuring significance

Although posterior distributions contain all the information about the parameters in the model, we use the Local False Sign Rate (LFSR) proposed by (*24*) to summarize them in a compact and interpretable value to perform significance hypothesis testing. Intuitively, the LFSR measures confidence in the sign of the effect. We are particularly interested in evaluating when the effect **D** _*j*_ of a condition **A** on an isoform *j* is significantly different from 0. We thus compute the LFSR as:

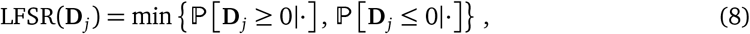

where ℙ [**D** _*j*_ ≥ 0| ·] and ℙ [**D** _*j*_ ≥ 0| ·] denote the posterior probabilities that **D** _*j*_ is greater or smaller than 0, respectively. We similarly define the LFSR for the other parameters.

### Simulation study

We simulate transcript, protein isoform, and peptide data under the two-layer linear model in Eq. 1. We simulate data in two different settings. In the first “easy” setting, we generate data without shared peptides. Specifically, we sample one transcript (*q* = 1) from a Normal distribution (i.e. **T**_*i j*_ ∼ 𝒩 (3, 1)), map it into a protein isoform **I**_*i j*_ with a random conversion weight (**W** _*j j*_ ∼ 𝒩 (1, 0.5)), add an effect due to a binary condition (**D** _*j*_ ∈ {0.33, 0.66, 1}), a random intercept term 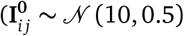, constant for all *i*) and random noise 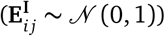, and then split it into two peptides (*r* = 2) using randomly assigned detectability scores 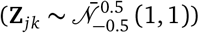 again adding random noise 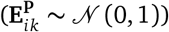. We define **A**_*i*_ = 1 for the first half of the sample, i.e. for *i* = 1, …, *n/*2, and **A**_*i*_ = 0 otherwise. In symbols:

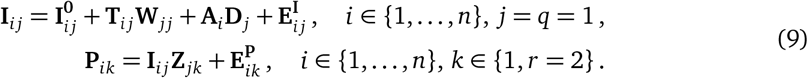

In the second “difficult” setting, we generate data that include shared peptides. Here, we sample five transcripts (*q* = 5) from a multivariate Normal distribution with identity covariance matrix (i.e. **T**_*i j*_ ∼ 𝒩 (3, 1)), map them to their corresponding protein isoforms **I**_*i j*_ with random conversion weights (**W** _*j j*_ ∼ 𝒩 (1, 0.5)), add an intercept term 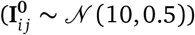, and introduce a binary condition effect (**D** _*j*_ ∈ {0.33, 0.66, 1}) to the first |**D**^*act*^ | ∈ {1, 2, 3} isoforms, i.e. **D** _*j*_ = **D** _*j*_ 𝟙 { *j* ≤|**D**^*act*^|}. The isoforms are then split into 10 peptides (*r* = 10) using random detectability scores, with *s* = 30% of the detectability matrix entries being nonzero, i.e. 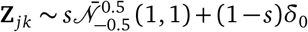. As before, we define **A**_*i*_ = 1 for the first half of the sample, i.e. for *i* = 1, …, *n/*2, and **A**_*i*_ = 0 otherwise. In symbols:

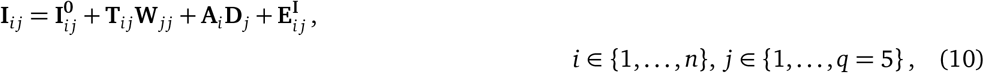

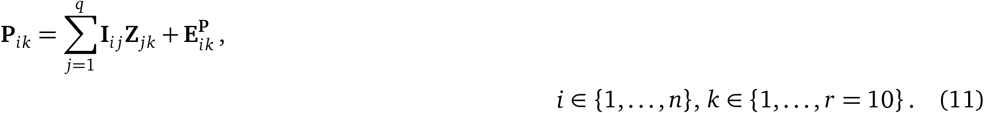

We run each simulation experiment with three different sample sizes (*n* = 100, 200, 500), and with 25 different seeds. We run `PAQu` for 3000 Gibbs sampling iterations, burning in the first 2000 iterations.

### Relationship to other methodology

We want to point out that, from a technical standpoint, our approach shares some similarities with (*47, 48*), as both employ a variant of Bayesian factor analysis. However, our method differs significantly in several key aspects. Specifically, we adopt different prior distributions, implement a distinct sampling algorithm, and target a different application focus. These differences result in a more tailored and efficient solution to the problem of protein isoform quantification, setting `PAQu` apart from previous approaches.

Moreover, by developing a tool for supervised factor analysis, our work has some connections to (*49–51*). However, these works only provide inference for latent factors (in our case, protein isoforms). Instead, `PAQu` provides estimates for a full class of parameters, shedding light on the biological processes behind protein formation.

#### Sample preparation, LC-MS*/*MS measurement, and database search for proteins

See In *SI Appendix*, Section C, for full details. In brief, brain samples were obtained from the University of Pittsburgh brain bank. A tissue sample of similar size was collected from each brain. Each sample was homogenized, its proteins were then solubilized, digested into peptides, TMT labeled, fractionated, and analyzed with an Orbitrap Tribrid Eclipse mass spectrometer. Samples were processed in batches of 14 samples, which will be described as a plex. RNA transcripts observed in the same brains (*25*), as measured by RNA-seq, were used to inform which isoforms were likely to be found in these samples (isoform matching).

#### Data preprocessing

##### Peptide abundance data

We first normalize raw expression values of samples to account for differences in total expression levels across samples (see *SI Appendix*, Section C, for details). After log_2_-transforming normalized values, effects of batch (aka, plex), diagnosis, age, and postmortem interval are modeled using ordinary least squares (OLS) regression. Effects of plex were subtracted from the data, leaving residuals that capture biological variation.

##### Transcript expression data

Transcript expression data were produced by the CommonMind Consortium (*25*) and were downloaded from https://www.synapse.org, specifically syn29442530, which was accessed on February 4, 2025. (These data have been moved to the NIMH Data Archive.) We first normalize transcript expression counts, transform them to counts per million, set them on a log_2_ scale, and remove the effect of technical covariates (see *SI Appendix*, Section C, for details).

### PAQu analysis of schizophrenia dataset

We exploit the block-diagonal structure in **Z** to divide the original problem into 5241 independent blocks. We then combined isoforms represented by the same set of peptides into isoform groups. This reduces the number of isoforms from 9169 to 7271. Of these, 5831 are singleton isoforms while 3338 isoforms were grouped into 1440 isoform groups with 2 or more isoforms. Out of the 1440 isoform groups, 1396 represented isoforms from a single gene, the remaining 44 represent multiple genes. From here on, we will refer to the combined set of singleton isoforms and isoform groups as isoforms.

### GO enrichment analysis

We perform GO enrichment analysis using the `g:GOSt` tool for functional profiling contained in the `g:profiler` toolkit (*52*), version *e*111_*eg*58_*p*18_ *f* 463989*d*, with Benjamini-Hochberg significance threshold set at 0.05. All ENSG isoforms employed in the analysis are treated as background; isoforms with LFSR ≤ 0.05 are treated as foreground.

### Analysis of transcript and complementary isoform correlations

Of the 7271 isoforms, we selected for analysis those isoforms with low variability of estimated abundance across MCMC chains. Specifically, for each isoform and MCMC chain, we computed the mean abundance over 103 samples. Next, we computed the variance of the mean over the 10 MCMC chains. If the variance fell below one, the isoform was retained (*N* = 7107); otherwise, it was removed (*N* = 164). We then calculated the Pearson correlation, over samples, for each of the 7107 transcript/isoform pairs. To predict the observed correlation of transcript/isoform pairs, we estimated the effects for diagnosis and age from the estimated isoform abundances and adjusted transcript expression. It is well known that gene expression and protein abundance can vary by genotype. To quantify the impact of genetic variation, we gathered information on 26,134 eGenes for the cortex from the GTEx website (*53*). (By definition, an eGene is the genetic variant that explains the greatest variance in gene expression.) From this set, we selected 16,491 protein-coding genes and recorded the estimate of the slope (b) and allele frequency (f). The estimated variance explained by each eGene was then determined as 2 *f* (1 − *f*)*b*^2^, and this value was used to predict the observed correlation of transcript/isoform pairs. Transcript/isoform groups mapping to multiple genes were excluded from this analysis.

## Acknowledgements

L.T. wishes to thank Will Townes for useful feedback. This work was funded, in part, by grants from the NIMH (MH125235, MH123184) and the Simons Foundation (SFARI SF1018804). The Genotype-Tissue Expression (GTEx) Project was supported by the Common Fund of the Office of the Director of the National Institutes of Health, and by NCI, NHGRI, NHLBI, NIDA, NIMH, and NINDS. The transcript data used for the analyses described in this manuscript were obtained from the Synapse Portal (Sage Bionetworks). We thank the patients and families who donated material for these studies. We thank T. Lehner for his early and inspirational ideas about this project, as well as organizational and intellectual support. Data were generated as part of the CommonMind Consortium supported by funding from Takeda Pharmaceuticals Company Limited, F. Hoffmann-La Roche Ltd and NIH grants U01MH116442, R01MH085542, R01MH093725, P50MH066392, P50MH080405, R01MH097276, RO1-MH-075916, P50M096891, P50MH084053S1, R37MH057881, AG02219, AG05138, MH06692, R01MH110921, R01MH109677, R01MH109897, U01MH103392, and contract HHSN271201300031C through IRP NIMH. Brain tissue for the study was obtained from the following brain bank collections: the Mount Sinai NIH Brain and Tissue Repository, the University of Pennsylvania Alzheimer’s Disease Core Center, the University of Pittsburgh NeuroBioBank and Brain and Tissue Repositories, and the NIMH Human Brain Collection Core. CMC Leadership: Panos Roussos, Joseph Buxbaum, Andrew Chess, Schahram Akbarian, Vahram Haroutunian (Icahn School of Medicine at Mount Sinai), Bernie Devlin, David Lewis (University of Pittsburgh), Raquel Gur, Chang-Gyu Hahn (University of Pennsylvania), Enrico Domenici (University of Trento), Mette A. Peters, Solveig Sieberts (Sage Bionetworks), Thomas Lehner, Stefano Marenco, Barbara K. Lipska (NIMH). Rhesus Macaque tissue was provided by Scott Hemby through the Stanley Medical Research Institute for Funding for Non-Human Primate Research; and funded by NIMH grant R01MH074313. J.B. was supported in part by NARSAD Young Investigator Grant 27209 from the Brain & Behavior Research Foundation. G.E.H. was supported in part by NARSAD Young Investigator Grant 26313 from the Brain & Behavior Research Foundation. This work was supported in part through the computational resources and staff expertise provided by the Department of Scientific Computing at the Icahn School of Medicine at Mount Sinai.

## Supplementary Information

### A Main PAQu model specification

#### Priors

Here, we provide the complete list of prior distributions employed in our main model specification:

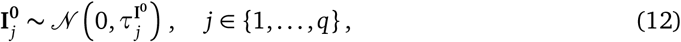

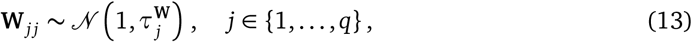

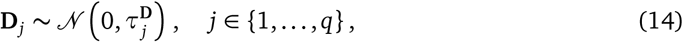

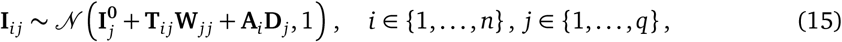

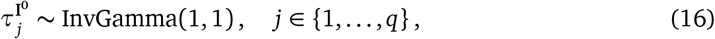

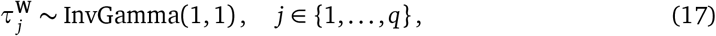

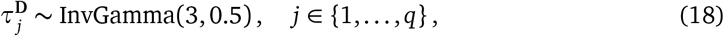

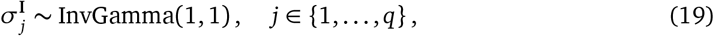

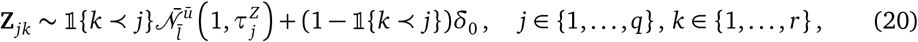

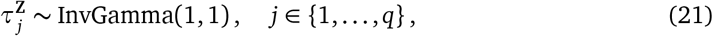

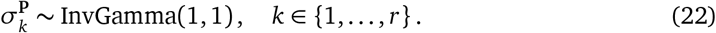

#### Posteriors

We exploit conjugacy in the Bayesian framework to obtain the following *full conditional* posteriors:

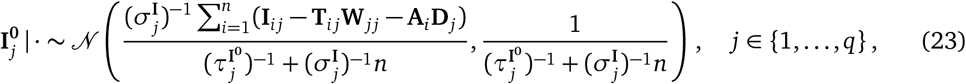

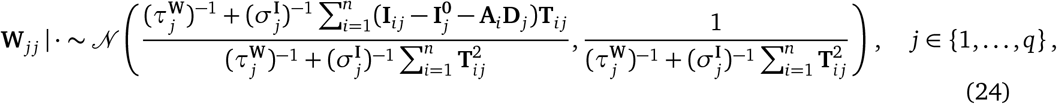

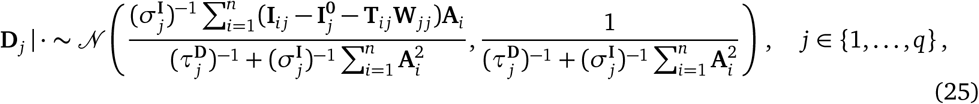

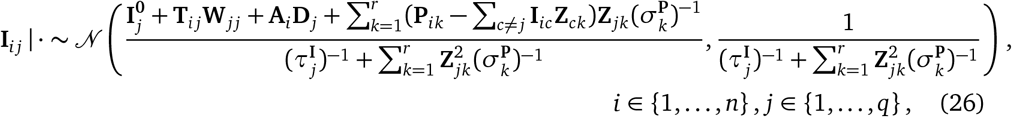

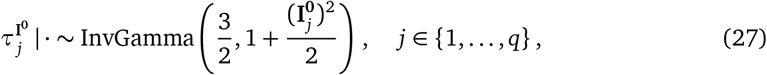

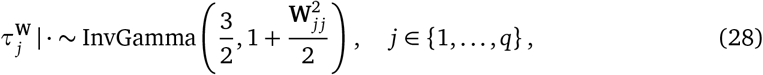

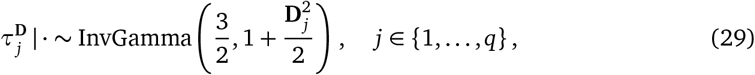

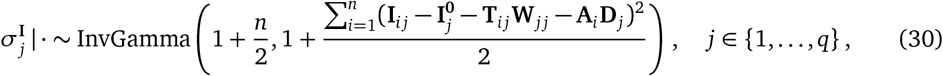

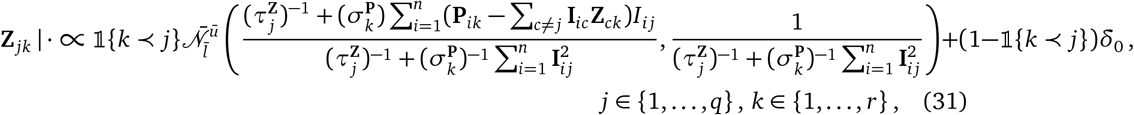

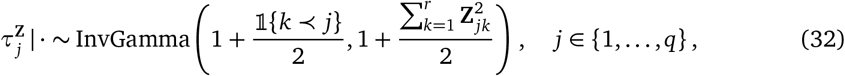

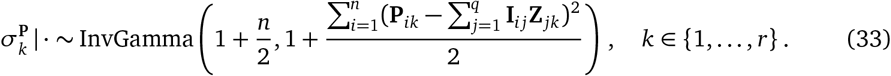

### Pseudocode and algorithmic implementation

#### Parameter initialization

We initialize **I** and **Z** from a truncated singular value decomposition (SVD) of **P**. Elements in **Z** are then masked according to the mask matrix **M**. We initialize **I**^**0**^, **D**, and **W** using ordinary least squares (OLS) estimates. In particular, we jointly regress each isoform on the corresponding transcript, the unit constant vector, and the condition vector **A**. All the other hyperparameters are originally initialized at 1.

#### Parallelization

By exploiting the sparsity in **Z**, we can divide the original problem into smaller, much easier-to-solve subproblems. In particular, **Z** presents a block-diagonal structure up to a permutation. Therefore, we only need to focus on one diagonal block at a time. Each diagonal block involves only the isoforms that share at least one peptide. In other words, isoforms with independent peptides can be analyzed separately. As a result, we can parallelize the computation block-by-block.

#### Pseudocode

Algorithm 1 below contains the pseudocode of `PAQu` running on an individual diagonal block.

##### Algorithm 1

`PAQu` algorithm pseudocode

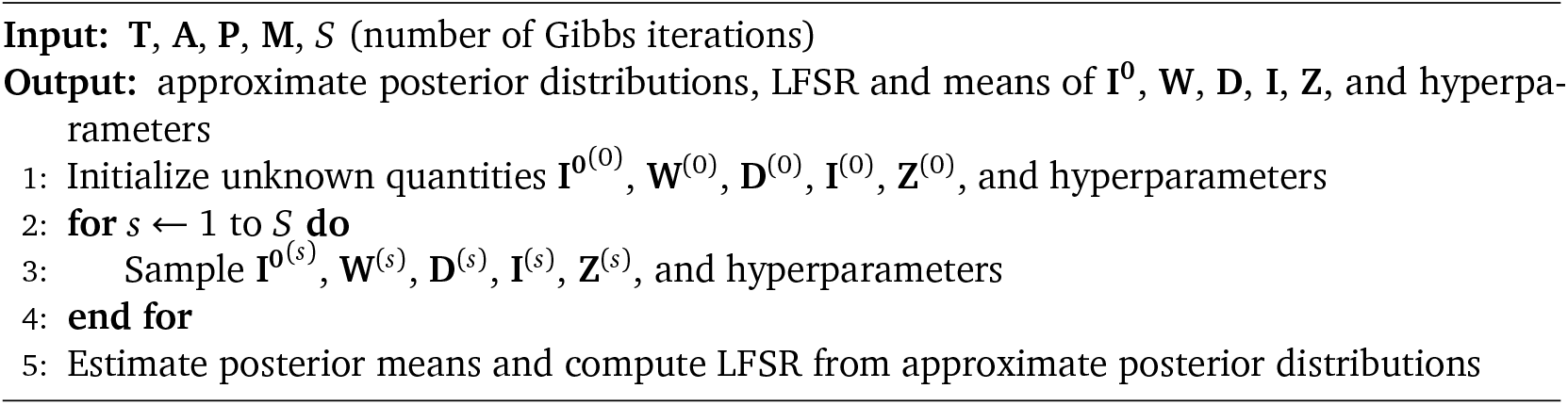

#### Spike-and-slab implementation

The prior structure of the problem changes only slightly. In particular, we need to introduce the following prior distributions:

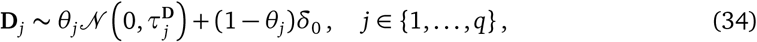

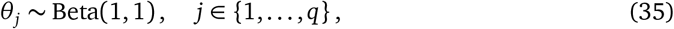

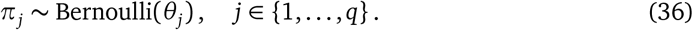

The derivation of the posterior distribution of *π* is not trivial. We first compute the log odds *y* (for numerical stability):

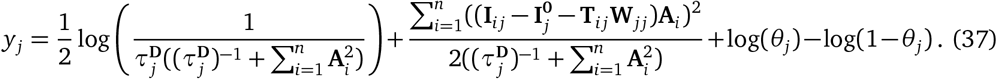

We then evaluate its logit transformation 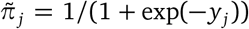. We thus sample from the posterior of *π*_*j*_ by computing 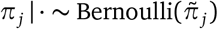.

The posterior distribution of *θ* is given by:

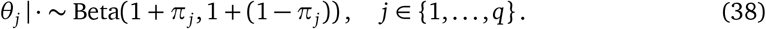

Finally, the posterior distribution of **D** _*j*_ is as before if *π*_*j*_ = 1; a Dirac mass at 0 otherwise.

### B Alternative PAQu model specifications

The minimal requirement for `PAQu` to run are the peptide abundance matrix and the mask matrix. Here, we describe several extensions of the model described in the main manuscript.

#### Absent information on transcripts or condition

In some scenarios, one may not have access to information on transcript expression levels or binary conditions. In these cases, `PAQu` operates by setting **W** = 0 and **D** = 0, respectively. Notice that, if information on transcripts is absent, then the user has to specify the number of protein isoform factors to employ to perform factor analysis. Therefore, in this case, *q* must be set in advance.

#### Limited information on transcripts

In some experiments, transcript expression levels may not have been measured, but the scientist has access to average expression levels from another study. In this case, these may be used as prior means for the intercept **I**^**0**^. In particular, define the average expression level for transcript *j* as *µ*_*j*_. Then the prior distribution for **I**^**0**^ is:

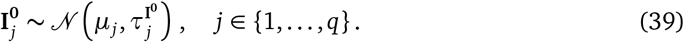

#### External covariates

In other settings, one may have access to additional information about protein isoforms or peptides. `PAQu` can accomodate external covariates in a straightforward manner. First, define **X**^**I**^ and **X**^**P**^ as the *n × p*^*I*^ and *n × p*^*P*^ matrices of external covariates that provide information on protein isoforms and peptides, respectively. We incorporate them into the model in Eq. 1:

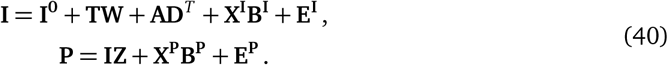

Here, the matrices **B**^**I**^ and **B**^**P**^ are *p*^*I*^ *× q* and *p*^*P*^ *× r*, capturing the effects of external covariates **X**^**I**^ and **X**^**P**^ on isoforms and peptides. The interpretation of the other variables is the same as before.

In this new setting, we need to define the following prior distributions:

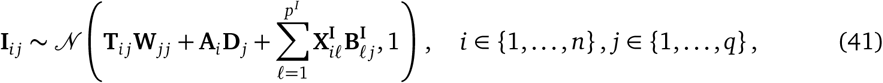

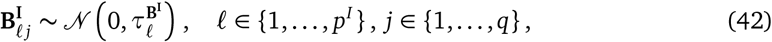

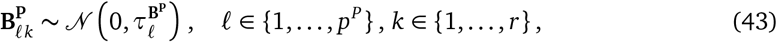

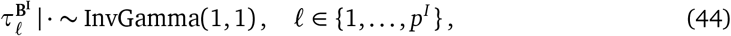

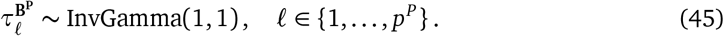

By similar computations as above, we get the following posteriors:

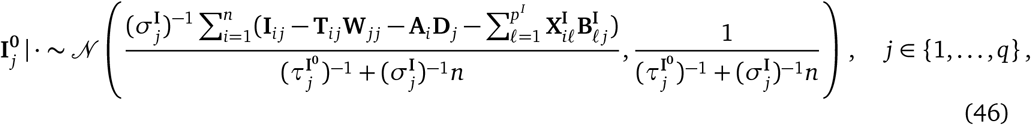

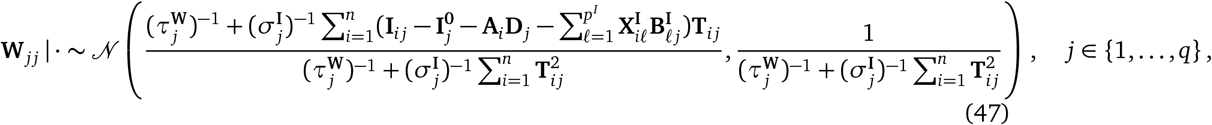

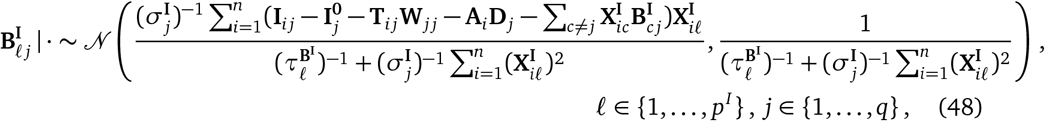

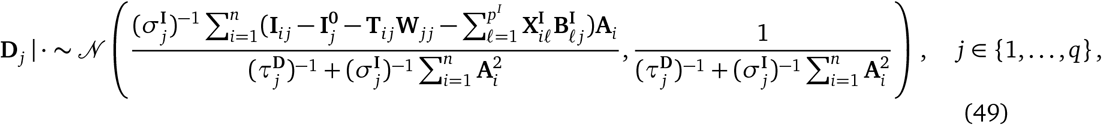

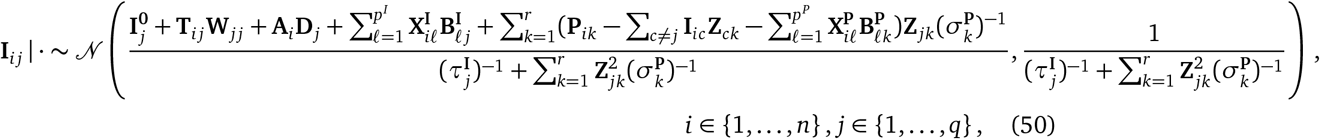

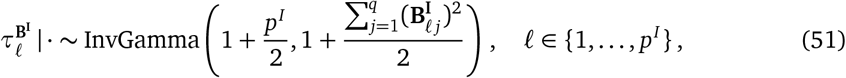

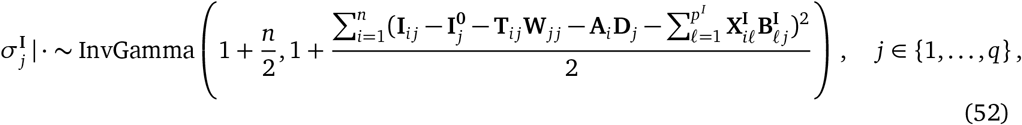

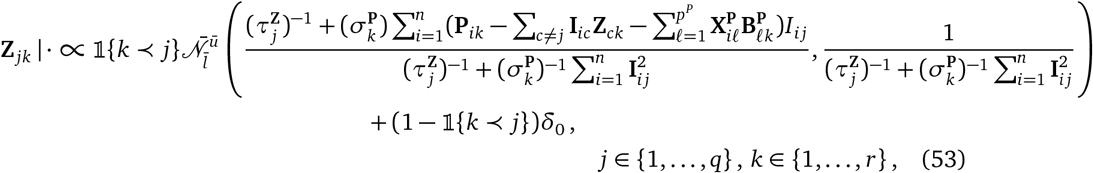

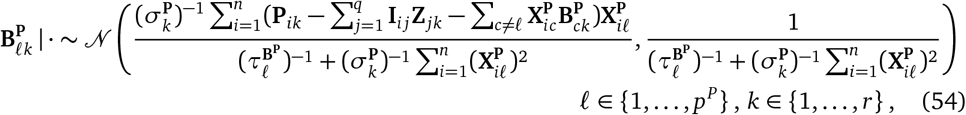

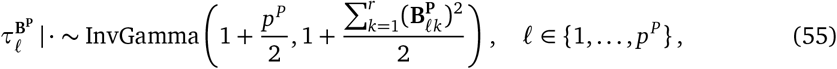

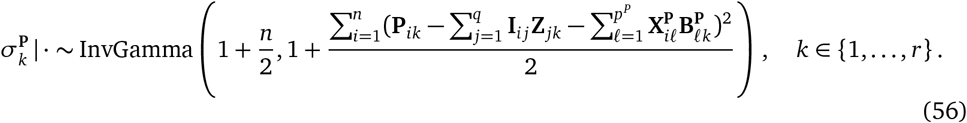

#### Collapsed isoforms

In some instances where two or more isoforms share exactly the same peptides and transcript expression levels do not bring any relevant prior information, `PAQu` may provide unstable estimates across replicate chains. Therefore, we also implement a *collapsed isoforms* strategy where we estimate the abundance of identical isoforms as they were a single meta-isoform. We take the average of their transcript expression levels as prior, if available. This allows `PAQu` to provide stable estimates of meta-isoforms abundance levels, which can still carry valuable biological insights.

### C Isoform and Transcript Characterization in Brain

#### Processing of Protein

##### Human tissue sample collection and preparation

Human brain specimens were obtained during autopsies conducted at the Allegheny County Office of the Medical Examiner after receiving consent from the next-of-kin. All procedures were approved by the Committee for the Oversight of Research and Clinical Training Involving Decedents and the Institutional Review Board for Biomedical Research, University of Pittsburgh, Pittsburgh, Pennsylvania. A subset of previously characterized Sz (n = 56) and control subjects (n = 56) were selected (*25*). When possible, subjects were organized into pairs matched by age, sex, and PMI. An independent panel of experienced clinicians made consensus diagnoses foloowing Diagnostic and Statistical Manual of Mental Disorders Fourth Edition (DSM-IV) or their absence using a previously described method. For area identification and blocking, grey matter was collected from the dACC by taking 40*µm* sections and stored frozen at -80°C. Experimenters were blinded to case-condition during sample preparation. SZ and CT pairs were kept together during all sample preparation steps. Samples were randomized and distributed in a block design for preparation and mass spectrometry analysis.

20mg of dACC grey matter was added to 10% SDS with 1x Protease (0.2%) and Phosphatase (0.5%) Inhibitors in a 1.5mL tube with 20mg of 0.2mm stainless steel beads. Samples were homogenized in a Bullet Blender Homogenizer (Next Advance) at speed 8 for 3 minutes, then supernatant diluted to 5x with (10%) SDS with 1x Protease (0.2%) and Phosphatase (0.5%) Inhibitors (Sigma, Cat. No., I3786, P5726, P0044) and frozen. Total protein concentration was assessed by micro-BCA (Thermo Fisher, Cat. No., 23235).

S-trap micro (Protifi, Cat. No. C02-micro-80) was used to digest proteins into tryptic peptides. For each sample, 100 µg of total protein was diluted to 50 µl in 5% SDS, 50 mM TEAB buffer, reduced with 5.6 µl 200 mM dithiothreitol (DTT) at 95^◦^C for 10 min, cooled down for 10 min at room temperature (RT), and alkylated with 5.6 µl 400 mM iodoacetamide (IAA) at RT for 30 min. Samples were then acidified with 8.6 µl 12% phosphoric acid, mixed with 569 µl loading/wash buffer (90% methanol, 100 mM TEAB pH 7.1), and loaded into S-trap 96-well plate columns by centrifugation at 1,500 ×*g*. Trapped proteins were washed with 200 µl wash buffer and centrifuged at 1,500 ×*g* for 1 min at 20^◦^C a total of three times. Proteins were then digested on-column with 10 µg trypsin in 125 µl 50 mM TEAB for 60 min at 47^◦^C. Peptides were collected by sequential centrifugation at 1,500 ×*g* for 2 min at 20^◦^C in 125 µl 50 mM TEAB, 125 µl 0.2% formic acid (FA), and 125 µl 50% acetonitrile (ACN). Eluted peptides were combined and dried in a vacuum concentrator.

A pooled control was made from 15 µg aliquots from each sample, which was then split into four aliquots for digestion on S-trap midi columns (Protifi). Samples were reduced with 27.7 µl 200 mM DTT at 95^◦^C for 10 min, cooled down for 10 min at RT, and alkylated with 27.7 µl 400 mM IAA at RT for 30 min. They were then acidified with 30.5 µl 12% phosphoric acid, mixed with 2016 µl loading/wash buffer, and loaded into S-trap columns by centrifugation at 4,000 ×*g*. Trapped proteins were washed with 600 µl wash buffer, centrifuged at 4,000 ×*g* for 1 min at 20^◦^C three times, and digested on-column with 25 µg trypsin in 125 µl 50 mM TEAB for 60 min at 47^◦^C. Peptides were collected by sequential centrifugation at 4,000 ×*g* for 1 min at 20^◦^C in 500 µl 50 mM TEAB, 500 µl 0.2% formic acid (FA), and 500 µl 50% acetonitrile (ACN). Eluted peptides were combined into one tube and dried in a vacuum concentrator.

##### TMT labeling and High pH reverse phase fractionation

Peptide digests were resuspended in 100 µl 100 mM TEAB and labeled with 40 µl of 10 µg/µl TMTpro reagent in ACN at room temperature (RT) for 1 hr, then quenched with 2.3 µl 5% hydroxylamine for 15 min. Pooled control peptides were split into two aliquots, each resuspended in 1300 µl 100 mM TEAB and labeled with 400 µl of 10 µg/µl TMTpro reagent in ACN at RT for 1 hr. The reaction was quenched with 92.79 µl 5% hydroxylamine for 15 min. A total of 100 µg from each sample was labeled with channels 1–16 in each plex (TMT channels 126–133N), and combined along with 100 µg pooled control peptides labeled with channels 17 and 18 (TMT 134C–135N), resulting in 16 samples and 2 pooled controls per plex. TMT-labeled peptide plexes were dried in a vacuum concentrator and fractionated by offline high-performance liquid chromatography (HPLC).

800 µg from each plex was resuspended in reagent A (4.5 mM ammonium formate, 2% ACN) and loaded onto a Zorbax Extend-C18 Rapid Resolution column (4.6 *×* 250 mm, 3.5 µm) on a Vanquish Flex system (Thermo Fisher). Samples were eluted with a 109 min gradient of mobile phase reagent B (4.5 mM ammonium formate, 90% ACN) at a flow rate of 0.800 mL/min. The gradient program was as follows: 0% B for 13 min, increasing to 16% B over 60 min, 40% B for 4 min, 44% B for 5 min, 60% B for 13 min, 99% B for 4 min, then decreasing to 0% B for the remaining 10 min. A total of 48 fractions were eluted and collected, then concatenated down to 24 using the following lineup: 1 and 25, 2 and 26, 3 and 27, 4 and 28, and so on. Fractions were dried in a vacuum concentrator and vialed in 100 µl of 93% Buffer A (0.1% formic acid in water) and 7% Buffer B (0.1% formic acid in acetonitrile).

##### Mass spectrometry and raw data processing

TMT-labeled peptide fractions were resuspended in 2% acetonitrile/0.1% formic acid and ∼1 µg was loaded onto a heated PepMap RSLC C18 2 µm, 100 Å, 75 µm × 50 cm column (Thermo Fisher, Cat. No. ES903) and eluted over a 180 min gradient. Sample eluate was electrosprayed (2,000 V) into a Thermo Scientific Orbitrap Eclipse mass spectrometer for analysis. MS1 spectra were acquired at a resolving power of 120,000. MS2 spectra were acquired in the ion trap with CID (35%) in centroid mode. Real-time search (max search time = 34 s; max missed cleavages = 1; Xcorr = 1; dCn = 0.1; ppm = 5) was used to select ions for synchronous precursor selection for MS3. MS3 spectra were acquired in the Orbitrap with HCD (60%) with an isolation window = 0.7 m/z and a resolving power of 60,000, and a max injection time of 400 ms. 4 µl (out of 20) of the TMT-labeled phosphopeptide enrichments were loaded onto a heated PepMap RSLC C18 2 µm, 100 Å, 75 µm × 50 cm column and eluted over a 180 min gradient: 1 min 2% B, 5 min 5% B, 160 min 25% B, 180 min 35% B. Sample eluate was electrosprayed (2,000 V) into a Thermo Scientific Orbitrap Eclipse mass spectrometer for analysis. MS1 spectra were acquired at a resolving power of 120,000. MS2 spectra were acquired in the Orbitrap with HCD (38%) in centroid mode with an isolation window = 0.4 m/z, a resolving power of 60,000, and a max injection time of 350 ms.

Raw MS files were processed in Proteome Discoverer version 2.4 (Thermo Fisher). MS spectra were searched against a custom FASTA database (see below). The SEQUEST search engine was used (enzyme = trypsin, max. missed cleavage = 3, min. peptide length = 6, precursor tolerance = 10 ppm). Static modifications included carbamidomethyl (C, +57.021 Da), and TMT labeling (N-term and K, +304.207 Da for TMTpro16). Dynamic modifications included oxidation (M, +15.995 Da), phosphorylation (S, T, Y, +79.966 Da, only for the phosphopeptide dataset), acetylation (N-term, +42.011 Da), Met-loss (N-term, − 131.040 Da), and Met-loss + acetyl (N-term, − 89.030 Da). Peptide spectral matches (PSMs) were filtered by the Percolator node (max Delta Cn = 0.05, target FDR (strict) = 0.01, and target FDR (relaxed) = 0.05). Proteins were identified with a minimum of 1 unique peptide and protein-level combined *q*-values < 0.05. Reporter ion quantification was based on intensity values with the following settings: integration tolerance = 20 ppm, method = most confident centroid, co-isolation threshold = 70, and SPS mass matches = 65.

##### Construction of brain region and cohort-specific FASTA file

The RNAseq database was queried for transcripts expressed at levels of ≥ 3 counts-per-million in ≥ 75% of the 103 samples, which yielded 23,185 transcripts meeting this criteria. These sequences were then transformed into a cohort- and brain area-specific protein sequence database using a 3-frame translation. We then compared searches with the custom FASTA to standard databases (e.g., Swiss-Prot) and supplemented our list with the single canonical “protein” sequence from any gene not represented in the RNAseq dataset.

### D Analysis of Isoform/Transcript Abundances

For the peptide dataset, we generated peptide measurements from samples of the anterior cingulate cortex (ACC) from brains preserved by the University of Pittsburgh brainbank. A tissue sample was taken from each of 56 individuals diagnosed with schizophrenia (case) and 56 unaffected individuals whose samples acted as controls. There were 46 matched case-control pairs (SEX, AGE, and PMI), which were always in the same plate and plex, and 10 cases and 10 controls that were evaluated as pairs, and they were similar in age and PMI, but not necessarily for sex. The samples were distributed over 8 plates; in addition to the 112 samples, 20 pooled samples (pools created from equal parts of all samples) were evaluated. Of the original 252,412 identified peptides, some had no measured abundance (46,214), and they were removed; likewise, peptides that could not be mapped to a protein (531) or peptides that had no detected abundance for at least half the samples and pools were removed (119,608). Remaining were 86,059 peptides for further analysis.

To impute missing values, starting values were obtained through the R package softImpute (*54*) using the pooled samples as a reference. After this, a variational autoencoder (scVAEIT) (*55*) was applied only to the data from samples and thereby the final imputed values were obtained. Covariates plex, diagnosis, sex, ancestry, age, pmi, and pH (adjusted for the effect of DX, i.e., residual pH) were included as predictor variables in VAEIT’s imputation process.

**Figure 6:**
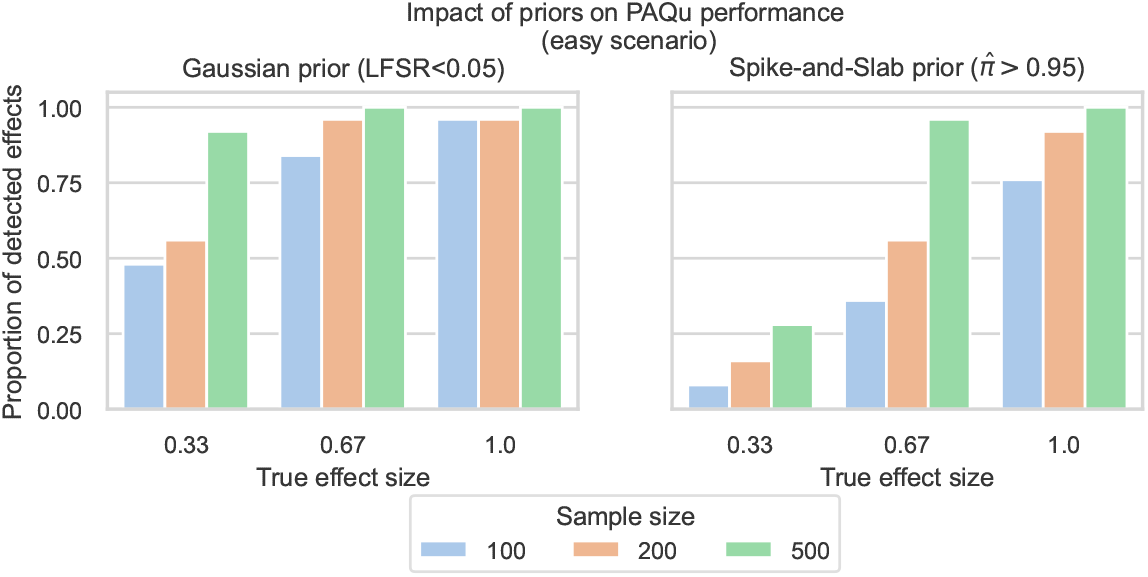
Simulation results, easy scenario, Gaussian vs spike-and-slab.

The imputed abundances for samples were normalized using sample lane normalization and then log_2_-transformed. Abundances were fit to a model with both biological (diagnosis, age, and postmortem interval) and technical predictors (plex) using ordinary least squares. The original abundance data were then adjusted for the effect of plex and pmi. By this process, we generated adjusted values for both transcripts and isoforms that still included effects for diagnosis and age. The resulting residuals were then used in subsequent analyses.

Transcript expression data were produced by the CommonMind Consortium (*25*) and were downloaded from https://www.synapse.org, specifically syn29442530, accessed on February 4, 2025. Expression data were available from 473 samples. Subjects from whom samples were drawn were either diagnosed with schizophrenia or were unaffected (controls). These samples had count data for 207,749 transcripts. The observed expression counts were normalized using calcNormFactors from R package edgeR and subsequently counts per million (cpm) were calculated using cpm from the same R package. After this step, the data were reduced to 53,918 transcripts with at least 50% of the samples having cpm > 1. These expression data were then log_2_-transformed for further processing. Following (*56*), adjusted expression was calculated by fitting a model with both biological (diagnosis, sex, age, and RIN) and technical variables (intronic rate, interegenic rate, ribosomal RNA rate, and institution) using ordinary least squares. The original data expresssion data were then adjusted for the effect of the technical covariates, as well as sex and RIN. After obtaining the adjusted transcript expression, the data were limited to samples matching those from which peptide measurements were taken.

Across the two datasets, 103 samples were in common: the protein abundance dataset consisted of 86,059 peptides; the expression dataset consisted of 53,918 transcripts. After selecting peptides that could be mapped to transcripts, and vice versa, there were 9169 transcripts and 79,040 peptides remaining.

**Figure 7:**
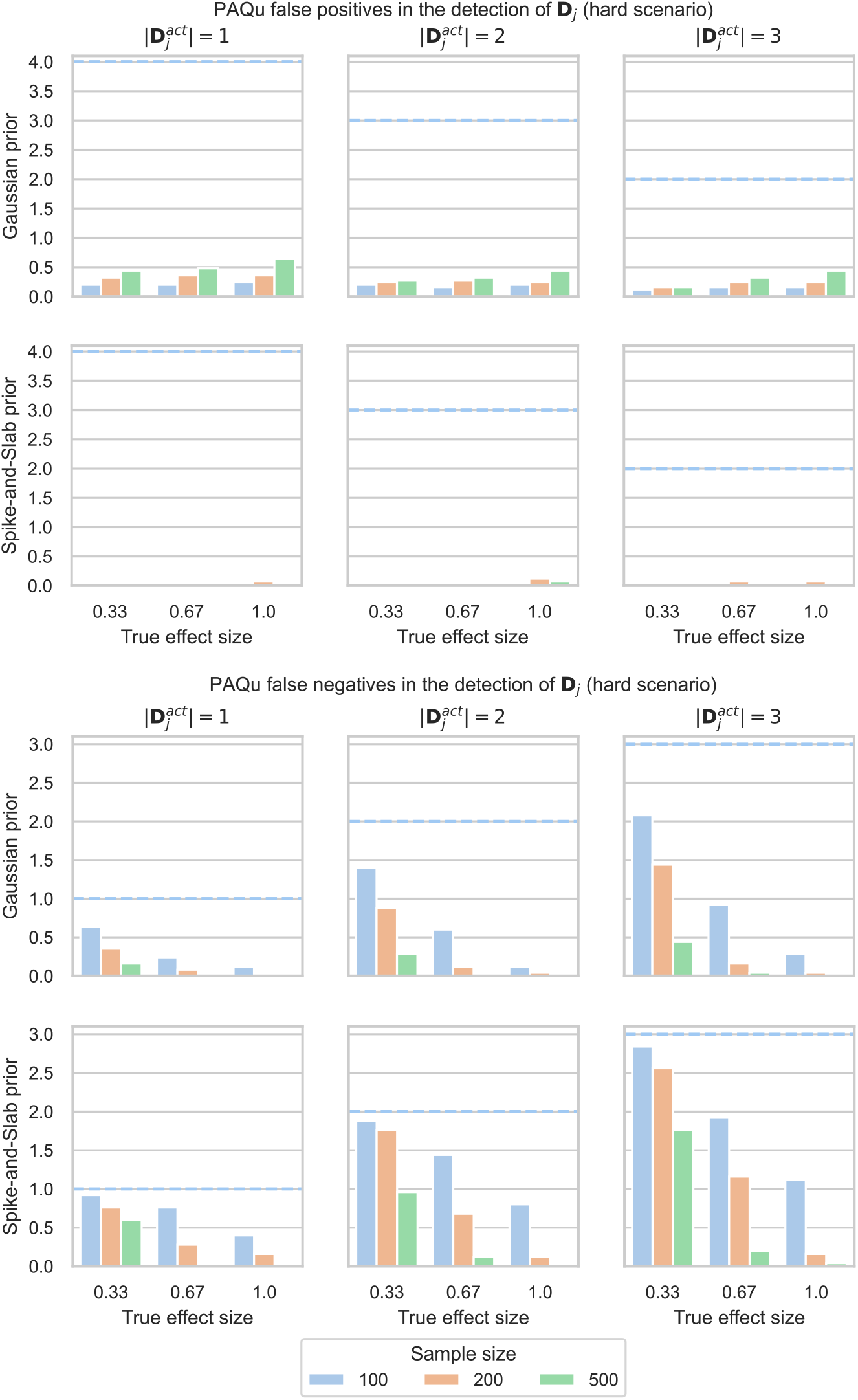
Simulation results, hard scenario, Gaussian vs spike-and-slab.

**Figure 8:**
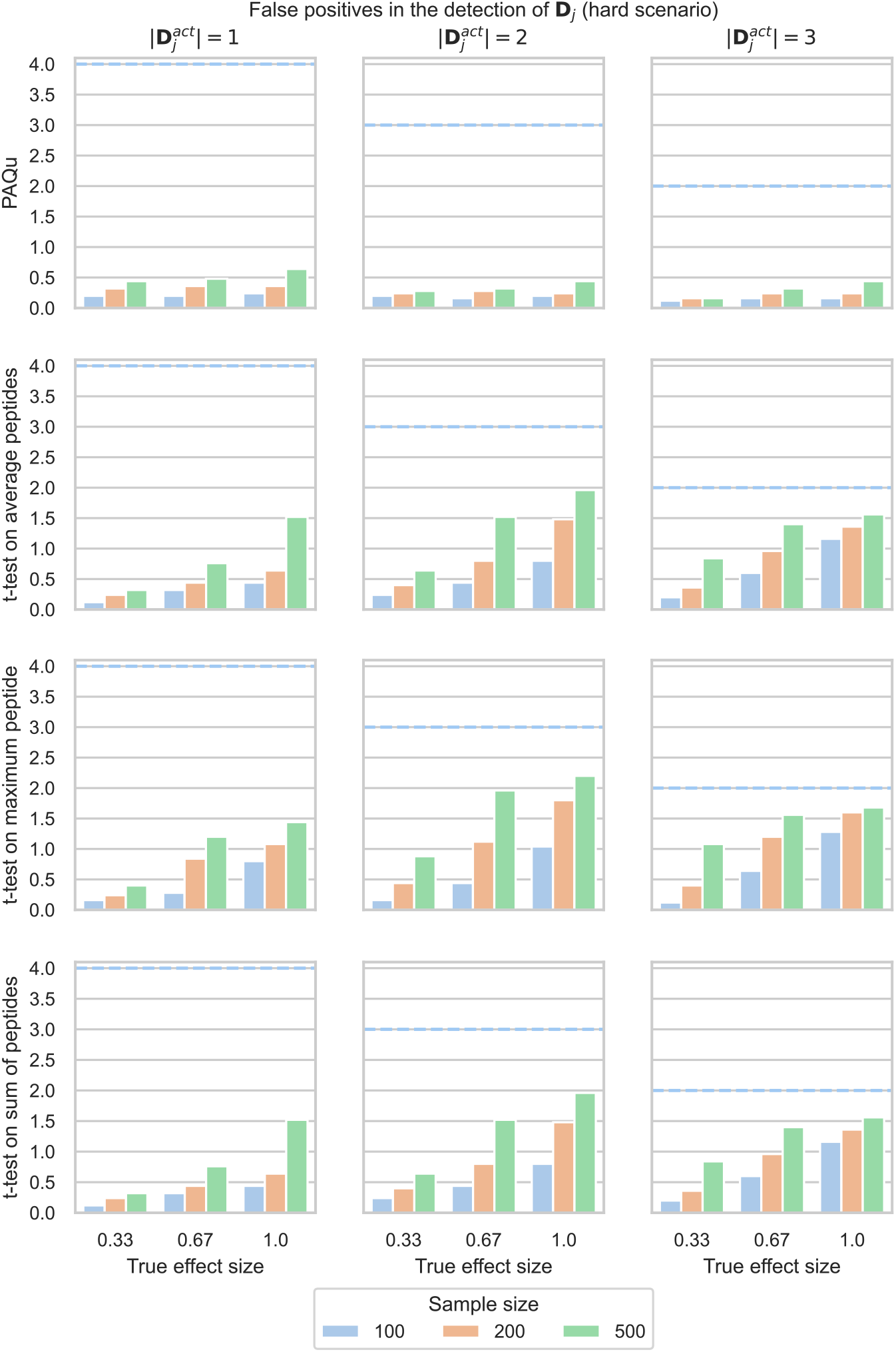
Simulation results, hard scenario, comparison. Blue dotted lines represent the maximum value the metric can take in any specific scenario.

**Figure 9:**
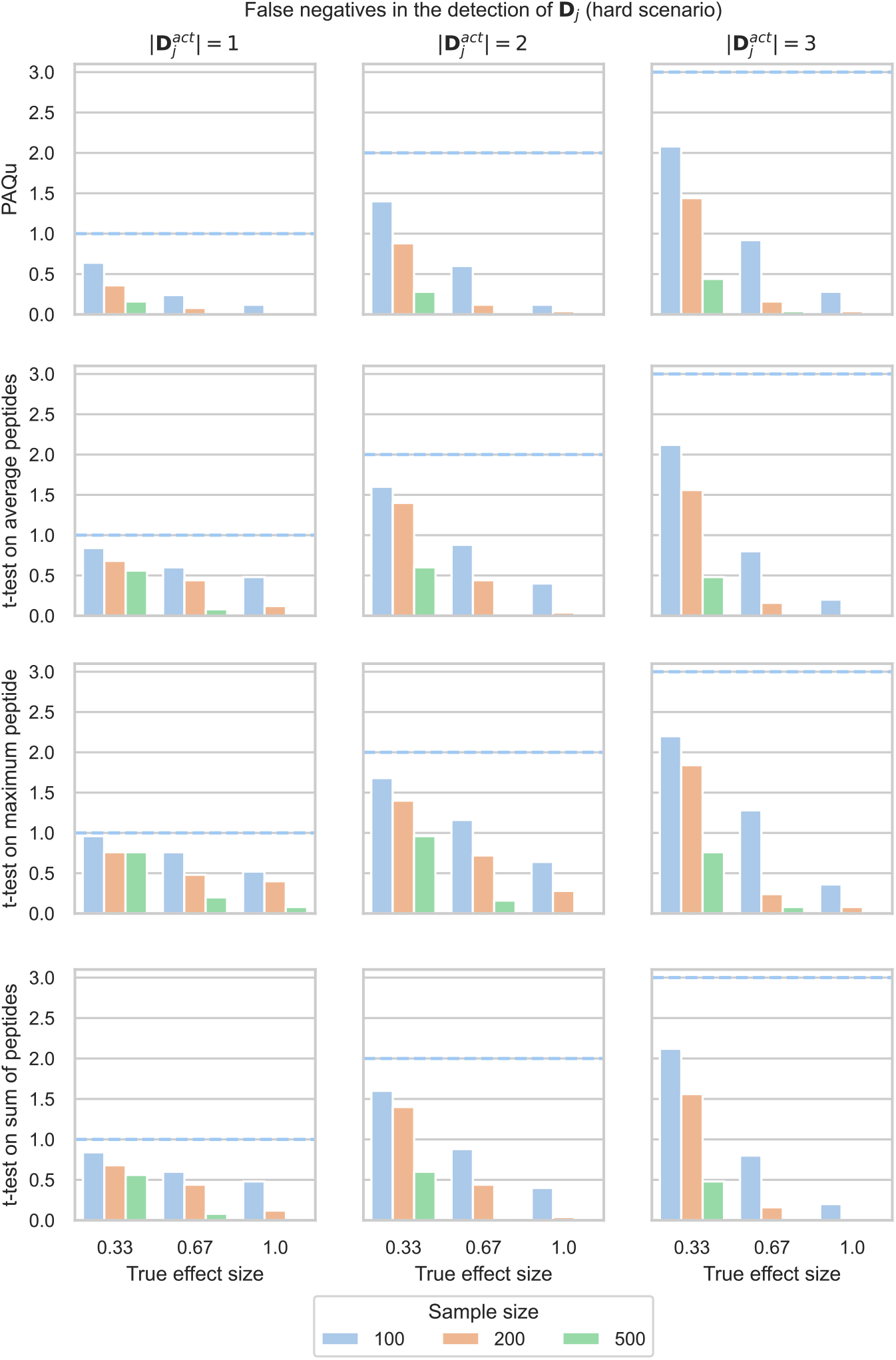
Simulation results, hard scenario, comparison. Blue dotted lines represent the maximum value the metric can take in any specific scenario.

**Figure 10:**
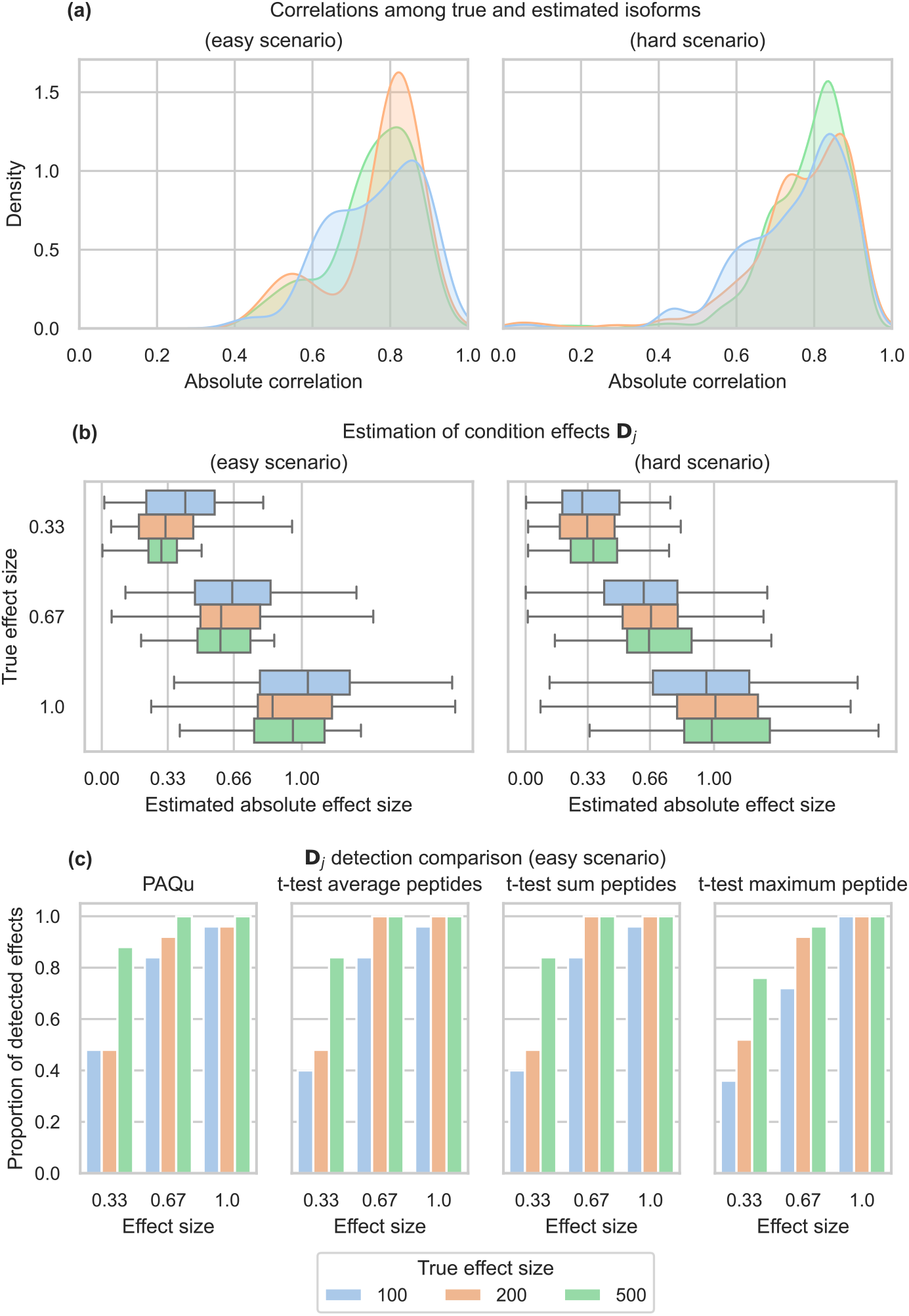
Simulation results, **W** = 0.

**Figure 11:**
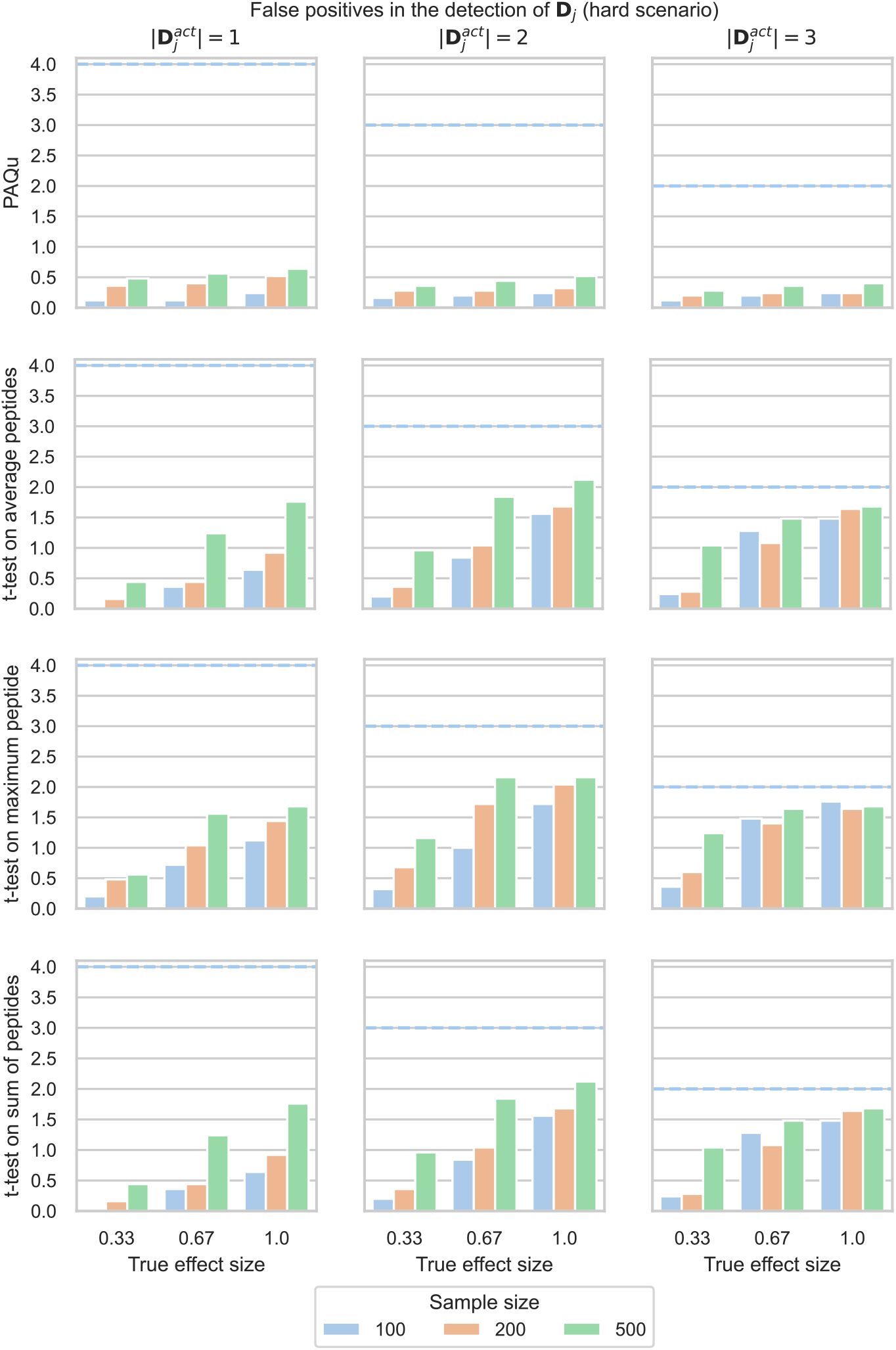
Simulation results, hard scenario, false positives, **W** = 0.

**Figure 12:**
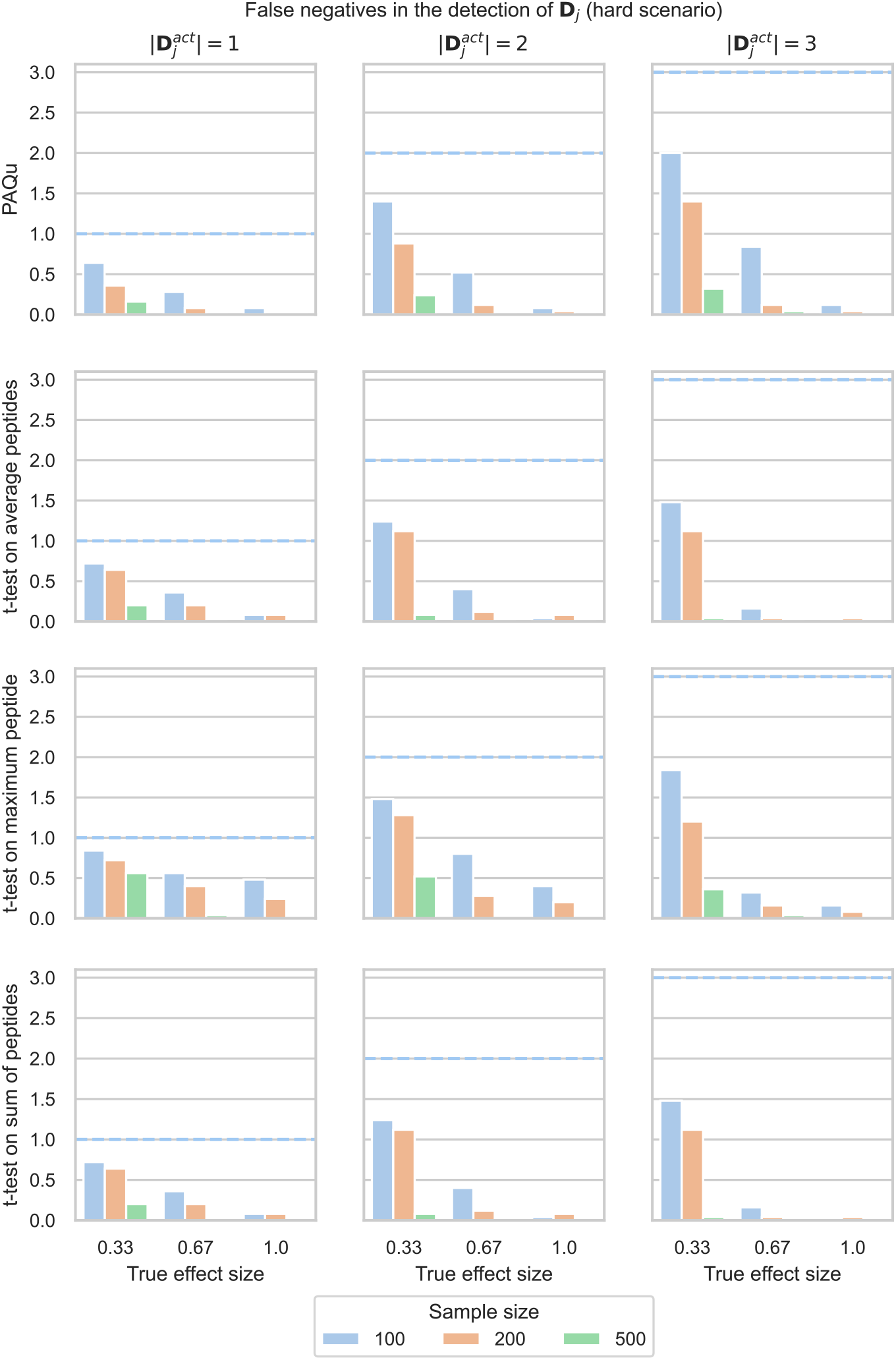
Simulation results, hard scenario, false negatives, **W** = 0.

**Figure 13:**
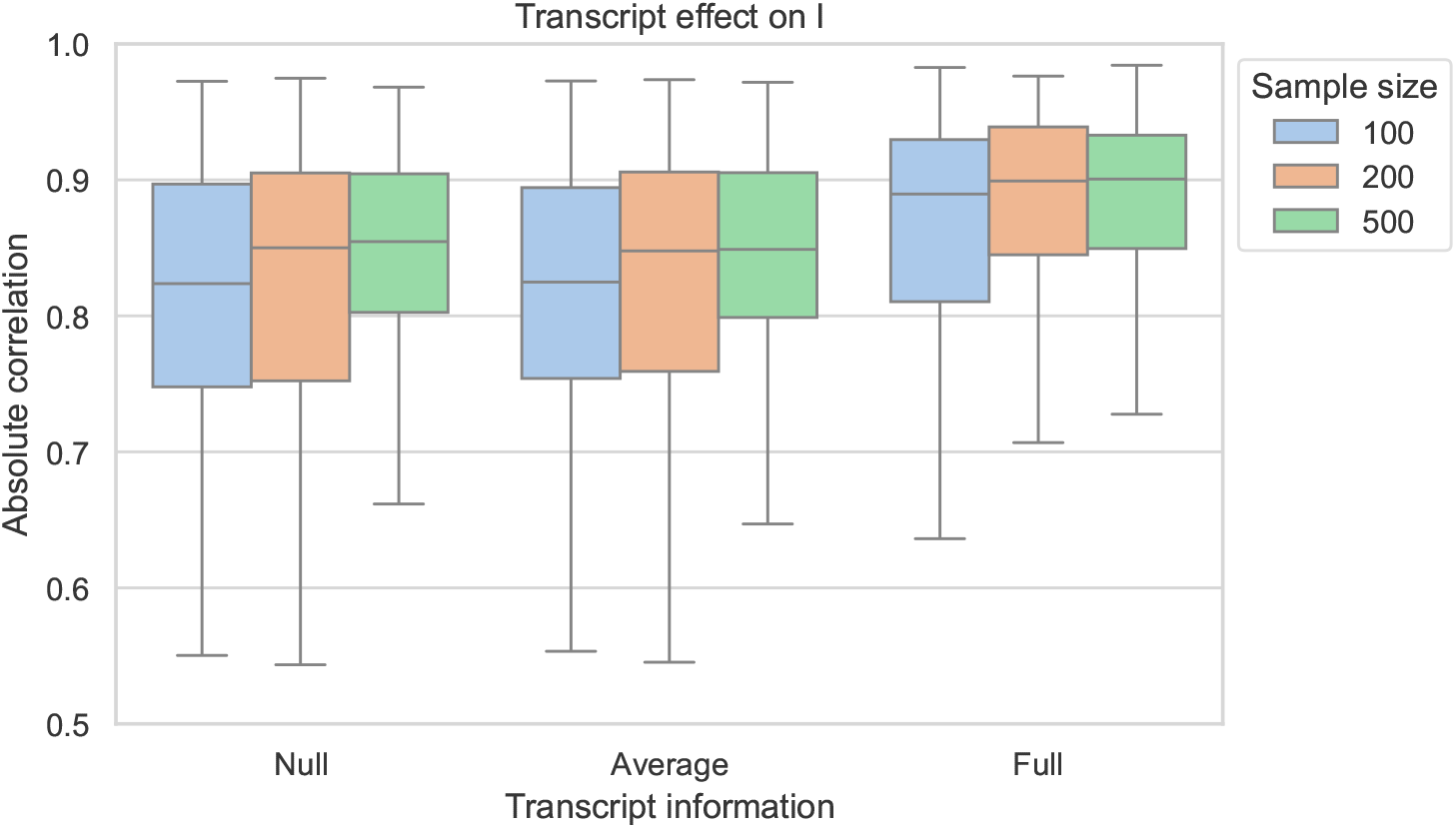
Effect on isoform abundance estimation of different availability of transcript information. |**D**^*act*^ | = 3, **D** _*j*_ = 0.33 for *j* = 1, 2, 3. Gaussian prior.

**Figure 14:**
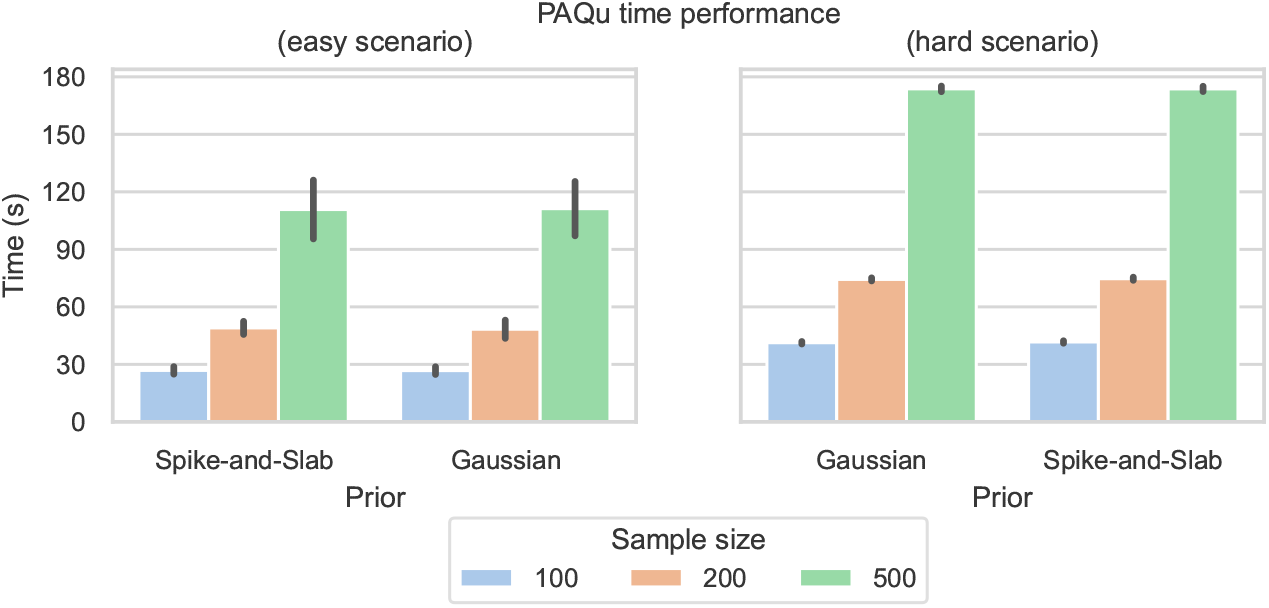
Simulation results, time.

**Figure 15:**
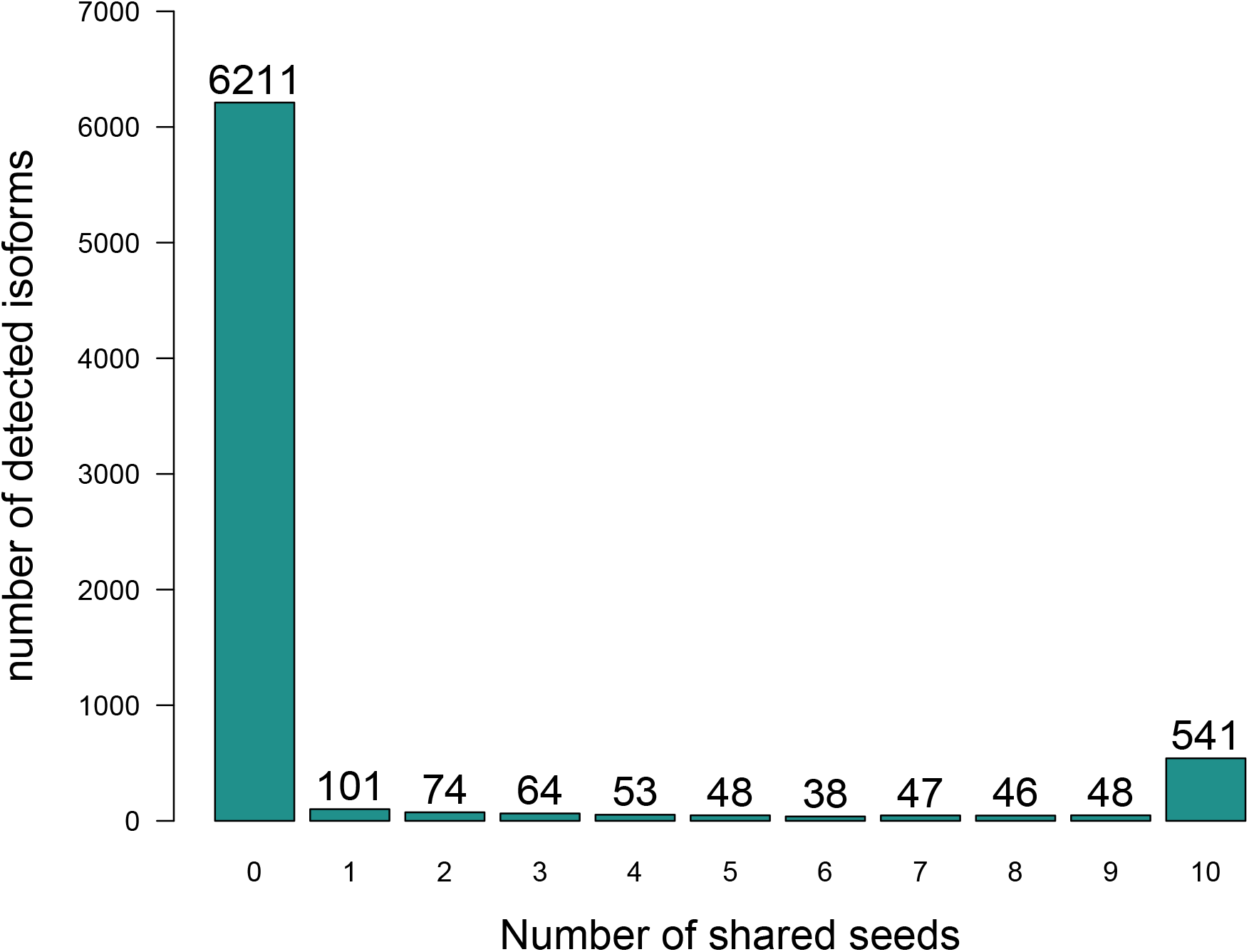
Isoform detection frequency as a function of the initial configuration seed

## References

(1) Kelemen, O., Convertini, P., Zhang, Z., Wen, Y., Shen, M., Falaleeva, M., and Stamm, S. (2013). Function of alternative splicing. Gene 514, 1–30.

(2) Lee, Y., and Rio, D. C. (2015). Mechanisms and Regulation of Alternative Pre-mRNA Splicing. Annu Rev Biochem 84, 291–323.

(3) Bradley, R. K., and Anczuków, O. (2023). RNA splicing dysregulation and the hallmarks of cancer. Nat Rev Cancer 23, 135–155.

(4) Zhang, Y., Qian, J., Gu, C., and Yang, Y. (2021). Alternative splicing and cancer: a systematic review. Signal Transduct Target Ther 6, 78.

(5) Li, D., McIntosh, C. S., Mastaglia, F. L., Wilton, S. D., and Aung-Htut, M. T. (2021). Neurodegenerative diseases: a hotbed for splicing defects and the potential therapies. Transl Neurodegener 10, 16.

(6) Gandal, M. J. et al. (2018). Transcriptome-wide isoform-level dysregulation in ASD, schizophrenia, and bipolar disorder. Science 362, DOI: 10.1126/science.aat8127.

(7) Wen, C. et al. (2024). Cross-ancestry atlas of gene, isoform, and splicing regulation in the developing human brain. Science 384, eadh0829.

(8) Sanders, S. J., Schwartz, G. B., and Farh, K. K.-H. (2020). Clinical impact of splicing in neurodevelopmental disorders. Genome Med 12, 36.

(9) Lesurf, R. et al. (2024). A validated heart-specific model for splice-disrupting variants in childhood heart disease. Genome Med 16, 119.

(10) Koko, M. et al. (2021). An identical-by-descent novel splice-donor variant in PRUNE1 causes a neurodevelopmental syndrome with prominent dystonia in two consanguineous Sudanese families. Ann Hum Genet 85, 186–195.

(11) Sinitcyn, P., Richards, A. L., Weatheritt, R. J., Brademan, D. R., Marx, H., Shishkova, E., Meyer, J. G., Hebert, A. S., Westphall, M. S., Blencowe, B. J., Cox, J., and Coon, J. J. (2023). Global detection of human variants and isoforms by deep proteome sequencing. Nat Biotechnol 41, 1776–1786.

(12) Miller, R. M. et al. (2022). Enhanced protein isoform characterization through long-read proteogenomics. Genome Biol 23, 69.

(13) Nesvizhskii, A. I., and Aebersold, R. (2005). Interpretation of shotgun proteomic data: the protein inference problem. Mol Cell Proteomics 4, 1419–40.

(14) Claassen, M., Reiter, L., Hengartner, M. O., Buhmann, J. M., and Aebersold, R. (2012). Generic comparison of protein inference engines. Mol Cell Proteomics 11, O110.007088.

(15) Meyer, J. G. (2021). Qualitative and Quantitative Shotgun Proteomics Data Analysis from Data-Dependent Acquisition Mass Spectrometry. Methods Mol Biol 2259, 297–308.

(16) Zhang, B., Chambers, M. C., and Tabb, D. L. (2007). Proteomic parsimony through bipartite graph analysis improves accuracy and transparency. J Proteome Res 6, 3549–57.

(17) Orsburn, B. C. (2021). Proteome discoverer—a community enhanced data processing suite for protein informatics. Proteomes 9, 15.

(18) Kong, A. T., Leprevost, F. V., Avtonomov, D. M., Mellacheruvu, D., and Nesvizhskii, A. I. (2017). MSFragger: ultrafast and comprehensive peptide identification in mass spectrometry– based proteomics. Nature methods 14, 513–520.

(19) Carlyle, B. C., Kitchen, R. R., Zhang, J., Wilson, R. S., Lam, T. T., Rozowsky, J. S., Williams, K. R., Sestan, N., Gerstein, M. B., and Nairn, A. C. (2018). Isoform-level interpretation of high-throughput proteomics data enabled by deep integration with RNA-seq. Journal of proteome research 17, 3431–3444.

(20) Miller, R. M., Jordan, B. T., Mehlferber, M. M., Jeffery, E. D., Chatzipantsiou, C., Kaur, S., Millikin, R. J., Dai, Y., Tiberi, S., Castaldi, P. J., et al. (2022). Enhanced protein isoform characterization through long-read proteogenomics. Genome biology 23, 69.

(21) Bollon, J., Shortreed, M. R., Jeffery, E., Jordan, B. T., Miller, R., Cavalli, A., Smith, L. M., Dewey, C. N., Sheynkman, G. M., and Tiberi, S. (2025). IsoBayes: a Bayesian approach for single-isoform proteomics inference. Bioinformatics, btaf450.

(22) Ishwaran, H., and Rao, J. S. (2005). Spike and slab variable selection: Frequentist and Bayesian strategies. The Annals of Statistics 33, 730–773.

(23) Hoff, P. D., A first course in Bayesian statistical methods; Springer: 2009; Vol. 580.

(24) Stephens, M. (2017). False discovery rates: a new deal. Biostatistics 18, 275–294.

(25) Fromer, M., Roussos, P., Sieberts, S. K., Johnson, J. S., Kavanagh, D. H., Perumal, T. M., Ruderfer, D. M., Oh, E. C., Topol, A., Shah, H. R., et al. (2016). Gene expression elucidates functional impact of polygenic risk for schizophrenia. Nature neuroscience 19, 1442–1453.

(26) Liu, Y., Beyer, A., and Aebersold, R. (2016). On the Dependency of Cellular Protein Levels on mRNA Abundance. Cell 165, 535–50.

(27) Cenik, C., Cenik, E. S., Byeon, G. W., Grubert, F., Candille, S. I., Spacek, D., Alsallakh, B., Tilgner, H., Araya, C. L., Tang, H., Ricci, E., and Snyder, M. P. (2015). Integrative analysis of RNA, translation, and protein levels reveals distinct regulatory variation across humans. Genome Res 25, 1610–21.

(28) Teo, G., Vogel, C., Ghosh, D., Kim, S., and Choi, H. (2014). PECA: a novel statistical tool for deconvoluting time-dependent gene expression regulation. J Proteome Res 13, 29–37.

(29) Lundberg, E., Fagerberg, L., Klevebring, D., Matic, I., Geiger, T., Cox, J., Algenäs, C., Lundeberg, J., Mann, M., and Uhlen, M. (2010). Defining the transcriptome and proteome in three functionally different human cell lines. Mol Syst Biol 6, 450.

(30) Jovanovic, M. et al. (2015). Immunogenetics. Dynamic profiling of the protein life cycle in response to pathogens. Science 347, 1259038.

(31) Bauernfeind, A. L., and Babbitt, C. C. (2017). The predictive nature of transcript expression levels on protein expression in adult human brain. BMC genomics 18, 322.

(32) Robins, C. et al. (2021). Genetic control of the human brain proteome. Am J Hum Genet 108, 400–410.

(33) GTEx Consortium et al. (2017). Genetic effects on gene expression across human tissues. Nature 550, 204–213.

(34) Ashburner, M. et al. (2000). Gene ontology: tool for the unification of biology. The Gene Ontology Consortium. Nat Genet 25, 25–9.

(35) Thomas, P. D., Ebert, D., Muruganujan, A., Mushayahama, T., Albou, L.-P., and Mi, H. (2022). PANTHER: Making genome-scale phylogenetics accessible to all. Protein Sci 31, 8–22.

(36) Gene Ontology Consortium et al. (2023). The Gene Ontology knowledgebase in 2023. Genetics 224, DOI: 10.1093/genetics/iyad031.

(37) Supek, F., Bošnjak, M., Škunca, N., and Šmuc, T. (2011). REVIGO summarizes and visualizes long lists of gene ontology terms. PloS one 6, e21800.

(38) Turrigiano, G. G. (2008). The self-tuning neuron: synaptic scaling of excitatory synapses. Cell 135, 422–435.

(39) Colameo, D., Rajman, M., Soutschek, M., Bicker, S., von Ziegler, L., Bohacek, J., Winterer, J., Germain, P.-L., Dieterich, C., and Schratt, G. (2021). Pervasive compartment-specific regulation of gene expression during homeostatic synaptic scaling. The EMBO Reports 22, EMBR202052094.

(40) Caterino, C., Ugolini, M., Durso, W., Jevdokimenko, K., Groth, M., Riege, K., Görlach, M., Fornasiero, E., Ori, A., Hoffmann, S., et al. (2025). Translational Remodeling of the Synaptic Proteome During Aging. Aging Cell 24, e70262.

(41) Seyfried, N. T., Dammer, E. B., Swarup, V., Nandakumar, D., Duong, D. M., Yin, L., Deng, Q., Nguyen, T., Hales, C. M., Wingo, T., et al. (2017). A multi-network approach identifies protein-specific co-expression in asymptomatic and symptomatic Alzheimer’s disease. Cell systems 4, 60–72.

(42) Chick, J. M., Kolippakkam, D., Nusinow, D. P., Zhai, B., Rad, R., Huttlin, E. L., and Gygi, S. P. (2015). A mass-tolerant database search identifies a large proportion of unassigned spectra in shotgun proteomics as modified peptides. Nat Biotechnol 33, 743–9.

(43) Sekar, A., Bialas, A. R., De Rivera, H., Davis, A., Hammond, T. R., Kamitaki, N., Tooley, K., Presumey, J., Baum, M., Van Doren, V., et al. (2016). Schizophrenia risk from complex variation of complement component 4. Nature 530, 177–183.

(44) Yilmaz, M., Yalcin, E., Presumey, J., Aw, E., Ma, M., Whelan, C. W., Stevens, B., McCarroll, S. A., and Carroll, M. C. (2021). Overexpression of schizophrenia susceptibility factor human complement C4A promotes excessive synaptic loss and behavioral changes in mice. Nature neuroscience 24, 214–224.

(45) de Sousa Nóbrega, I., Moysés, M. B. B., Yokota-Moreno, B. Y., Ferreira, G. G., and Sertié, A. L. (2025). Schizophrenia-associated complement component C4 induces maturation and reactivity in human astrocytes. Immunology Letters, 107065.

(46) Han, K. A., Jang, G., Lee, H.-Y., Kim, B., Kanato, T., Vas, V., Buday, L., Liu, X., Choi, S.-Y., Um, J. W., et al. (2025). CASKIN2 mediates PTP*σ*-orchestrated transsynaptic mechanisms at excitatory synapses. Proceedings of the National Academy of Sciences 122, e2509116122.

(47) Zhou, Y., Luo, K., Liang, L., Chen, M., and He, X. (2023). A new Bayesian factor analysis method improves detection of genes and biological processes affected by perturbations in single-cell CRISPR screening. Nature Methods 20, 1693–1703.

(48) Gygi, J. P., Konstorum, A., Pawar, S., Aron, E., Kleinstein, S. H., and Guan, L. (2024). A supervised Bayesian factor model for the identification of multi-omics signatures. Bioinformatics 40, btae202.

(49) Fan, J., Liao, Y., and Wang, W. (2016). Projected principal component analysis in factor models. Annals of statistics 44, 219.

(50) Li, G., Yang, D., Nobel, A. B., and Shen, H. (2016). Supervised singular value decomposition and its asymptotic properties. Journal of Multivariate Analysis 146, 7–17.

(51) Yu, S., Yu, K., Tresp, V., Kriegel, H.-P., and Wu, M. In Proceedings of the 12th ACM SIGKDD international conference on Knowledge discovery and data mining, 2006, pp 464–473.

(52) Kolberg, L., Raudvere, U., Kuzmin, I., Vilo, J., and Peterson, H. (2020). gprofiler2 – an R package for gene list functional enrichment analysis and namespace conversion toolset g:Profiler [version 2; peer review: 2 approved]. F1000Research 9.

(53) Lonsdale, J., Thomas, J., Salvatore, M., Phillips, R., Lo, E., Shad, S., Hasz, R., Walters, G., Garcia, F., Young, N., et al. (2013). The genotype-tissue expression (GTEx) project. Nature genetics 45, 580–585.

(54) Mazumder, R., Hastie, T., and Tibshirani, R. (2010). Spectral regularization algorithms for learning large incomplete matrices. The Journal of Machine Learning Research 11, 2287–2322.

(55) Du, J.-H., Cai, Z., and Roeder, K. (2022). Robust probabilistic modeling for single-cell multimodal mosaic integration and imputation via scVAEIT. Proceedings of the National Academy of Sciences 119, e2214414119.

(56) Hoffman, G., Ma, Y., Montgomery, K. S., Bendl, J., Jaiswal, M. K., Kozlenkov, A., Peters, M. A., Dracheva, S., Fullard, J. F., Chess, A., et al. (2022). Sex differences in the human brain transcriptome of cases with schizophrenia. Biological psychiatry 91, 92–101.

